# Extended synaptotagmin regulates plasma membrane-endoplasmic reticulum contact site structure and lipid transfer function *in vivo*

**DOI:** 10.1101/2019.12.12.874933

**Authors:** Vaisaly R Nath, Shirish Mishra, Bishal Basak, Deepti Trivedi, Padinjat Raghu

## Abstract

Inter-organelle communication between closely apposed membranes is proposed at Membrane Contact Sites (MCS). However the regulation of MCS structure and their functional relevance *in vivo* remain debated. The extended synaptotagmins (Esyt) are evolutionarily conserved proteins proposed to function at MCS. However, loss of all three Esyts in yeast or mammals shows minimal phenotypes questioning the functional importance of Esyt. We report that in *Drosophila* photoreceptors, MCS number is regulated by PLCβ activity. Photoreceptors of a null allele of *Drosophila* extended synaptotagmin *(dEsyt)* show loss of ER-PM MCS. Loss of dEsyt results in mislocalization of RDGB, an MCS localized lipid transfer protein, required for photoreceptor structure and function, ultimately leading to retinal degeneration. *dEsyt* depletion enhanced the retinal degeneration, reduced light responses and slower rates of plasma membrane PIP_2_ resynthesis seen in *rdgB* mutants. Thus, dEsyt function and PLCβ signaling regulate ER-PM MCS structure and lipid transfer in *Drosophila* photoreceptors.

## Introduction

One of the key attributes of eukaryotic cells is the existence of membrane bound organelles each with its own unique protein and lipid composition. The role of vesicular transport between organelles in establishing and maintaining this composition is well accepted. However it has also been observed that in intact cells, organelle membranes are often placed in close proximity to each other without undergoing fusion as described for vesicular transport. This concept of closely apposed organelle membranes is referred to as <u>*M*</u>embrane <u>*C*</u>ontact <u>*S*</u>ites (MCS) that are now defined as sites where organelle membranes are found separated by a gap of 10-30 nm. MCS were noted in the electron microscopy studies of George Palade (Palade and Porter, 1957) and subsequent studies have described MCS in several cell types (Carrasco and Meyer, 2014; Hayashi et al., 2008; Hepler et al., 1984; Suzuki and Hirosawa, 1994). The endoplasmic reticulum (ER), the largest organelle in the cell forms an elaborate tubular network that occupies much of the interior of the cell and therefore establishes MCS with most other organelles (Gatta and Levine, 2017) including the plasma membrane (PM). A number of molecular processes are proposed to occur at MCS; these include Ca^2+^ homeostasis, lipid transfer between organelle membranes and signaling interactions between proteins located at membranes on either side of the MCS [reviewed in (Saheki and De Camilli, 2017) (Balla et al., 2019)]. However, the mechanisms by which MCS are established and maintained as well as their importance for cell physiology *in vivo* remain unclear.

Every organelle has a unique lipid composition that is a critical determinant of its function. Since lipids are hydrophobic and cannot diffuse in aqueous cytosol, lipid composition depends on the transport of lipid to and away from organelle membranes. Such intracellular lipid trafficking is mediated both by vesicular transport as well as the activity of lipid transfer proteins (LTP). A major function that has been proposed for MCS is the activity of LTP at these sites and several studies have demonstrated that LTPs can be localized to membrane contact sites [reviewed in (Cockcroft and Raghu, 2018)]. Thus, the integrity of MCS is likely to be central to lipid homeostasis in eukaryotic cells. However, the depletion of multiple proteins localized to MCS and is thought to contribute to their structure and functions have showed limited effect on cell physiology, questioning the relevance and importance of MCS function for lipid homeostasis *in vivo*.

MCS between the PM and the ER are found in many cell types. Studies in yeast have shown that a major fraction of the ER shows a cortical localization (Pichler et al., 2001) and is confined in close proximity to the PM. The ER and PM have distinct lipid compositions; the ER is a major site of lipid biosynthesis and multiple lipids including phosphatidylinositol are enriched in this compartment. By contrast, the PM is enriched in distinct lipid classes including phosphatidylinositol 4, 5-bisphosphate (PIP_2_) and phosphatidylinositol 4-phosphate (PI4P) that are not found at the ER membrane. Remarkably, the PI4P phosphatase Sac1 is localized at the ER while its substrate PI4P is enriched at the PM. Elegant studies in yeast have proposed that Sac1 is localized to ER-PM MCS and functions to dephosphorylate PI4P in *trans* (Stefan et al., 2011). Proteomic studies in yeast have identified a set of 6 proteins, referred to as tethers that interact with Sac1 and are localized to ER-PM contact sites. These include Scs2 and Scs22 (VAP orthologs) (Loewen et al., 2007), Ist2 (member of TMEM16 ion channel family) (Manford et al., 2012) and the tricalbin proteins (Tcb1, Tcb2, Tcb3 ‒Esyt orthologs) (Manford et al., 2012). Depletion of all these proteins using a so-called *Δtether* strain, leads to the collapse of the cortical ER (cER) away from the PM and elevated PI4P levels at the PM (Manford et al., 2012). However, the *Δtether* strain shows remarkably mild phenotypes, with only a limited change in PM permeability described on heat stress (Omnus et al., 2016) raising the question of whether the presence of contact sites is required to support cell physiology.

Extended synaptotagmins (Esyts) are a group of ER anchored proteins belonging to the Tubular Lipid Binding (TULIP) superfamily that comprise an N-terminal transmembrane domain, a synaptotagmin like mitochondrial lipid binding protein domain (an SMP domain) and several acidic phospholipid binding C2 domains that bind to PM lipids (Chang et al., 2013). Mammalian cells have three independent genes that encode for Esyt: *Esyt1, Esyt2, and Esyt3*. Studies in mammalian cells have described localization of these proteins to ER-PM contact sites and proposed a role for Esyt in mediating lipid transfer function (Giordano et al., 2013). It has been proposed that Ca^2+^ and PIP_2_ mediate the localization and tethering of Esyt to ER-PM contact sites (Bian et al., 2018; Yu et al., 2016) and that this is linked to the lipid transfer activity of the SMP domain (Idevall-hagren et al., 2015; Yu et al., 2016). Thus, two independent roles have been proposed for Esyt; one as a member of a group of proteins that function as a tether at ER-PM contact sites and another as a lipid transfer protein via its SMP domain. Yet, the yeast *Δtether* strain, that lacks all Esyt genes shows remarkably mild phenotypes (Omnus et al., 2016). A triple knockout of mouse Esyt1, 2 and 3 is viable and shows no major changes in brain morphology, synaptic protein composition and stress response (Sclip et al., 2016). Even so, another *in vitro* study on *Esyt2*^−/−^*/Esyt3*^−/−^ mouse embryonic fibroblast cells exhibit declined migratory potential and are vulnerable to stressed culture conditions (Herdman et al., 2014) questioning the role of Esyts in supporting function *in vivo*. Interestingly, in *Drosophila*, Esyt is reported to be localized to the presynaptic compartment in the neuromuscular junction and implicated in regulating neurotransmitter release and synaptic growth (Kikuma et al., 2017). Therefore, the *in vivo* function of this highly conserved protein family and the role of Esyt in supporting contact site functions remain unresolved.

In *Drosophila* photoreceptors, sensory transduction is mediated by G-protein coupled phospholipase C mediated PIP_2_ turnover (Raghu et al., 2012). The biochemical reactions that constitute the PIP_2_ cycle in these cells is organized around an ER-PM contact site between the sub-microvillar cisternae (SMC), a specialized compartment of the ER and the microvillar PM [reviewed in (Yadav et al., 2016)]. Localized to this contact site is a multi-domain protein Retinal Degeneration B (RDGB), that includes an N-terminal phosphatidylinositol transfer domain (PITP) that can transfer PI and PA i*n vitro*; binding of PI to the PITP domain is essential for the *in vivo* function of RDGB (Yadav et al., 2015). Loss of this function in flies lacking RDGB protein results in key *in vivo* phenotypes: reduced electrical response to light (reduction in ERG amplitude), delay in PIP_2_ resynthesis and light-dependent degeneration of photoreceptors (retinal degeneration) [reviewed in (Cockcroft et al., 2016)]. Thus, *Drosophila* photoreceptors offer an ideal *in vivo* model to test the mechanisms that control ER-PM contact site structure and lipid transfer. In this study, we have generated a loss-of-function mutant for the only Esyt in *Drosophila* and studied its phenotypes in photoreceptors. We find that PLCβ dependent Esyt function is required to support ER-PM contact site structure as well as the lipid transfer function of RDGB *in vivo*.

## Results

### A single gene encodes for Esyt in *Drosophila*

Bioinformatic analysis revealed that the *Drosophila* genome encodes a single ortholog of human Esyt and the tricalbin proteins of yeast (Fig 1A). The *Drosophila* extended synaptotagmin gene (*dEsyt*) is CG6643 located on the third chromosome (cytolocation 96A7-96A7). *dEsyt* encodes for four different transcripts: RA, RB, RC and RD that are identical except for a 75 bp sequence between the 2^nd^ and 3^rd^ C2 domain coding regions where each isoform has different sequences. Sequence comparison for all four transcripts showed that the RA CDS is the shortest, lacking 100 nucleotides at the 5’ end and the RD transcript specifically lacks 9 nucleotides which encode for 3 amino acid residues that are identical in the other three transcripts. When aligned with human Esyt1 (GI: 296317244), CG6643 shows 93% query coverage and 35% sequence identity with all four isoforms. The protein domain structure of dEsyt is conserved with that of the human and yeast orthologs (Fig 1B). For e.g in hEsyt proteins, a single N-terminal SMP domain and multiple C2 domains are described. dEsyt sequence shows the presence of an SMP domain and three C2 domains. This feature along with the conserved interdomain distance makes dEsyt most similar to hEsyt2 and hEsyt3. Transcriptome analysis indicates that *dEsyt* is ubiquitously expressed across tissues and during all developmental stages (http://flybase.org/reports/FBgn0266758). However q-PCR analysis showed that *dEsyt* transcripts are enriched in the head compared to the body (Fig 1C); however a comparison between transcript expression in heads of wild type and *so*^*D*^ (that lacks eyes) suggests that it is not enriched in the eye (Fig 1D).

**Figure 1:**
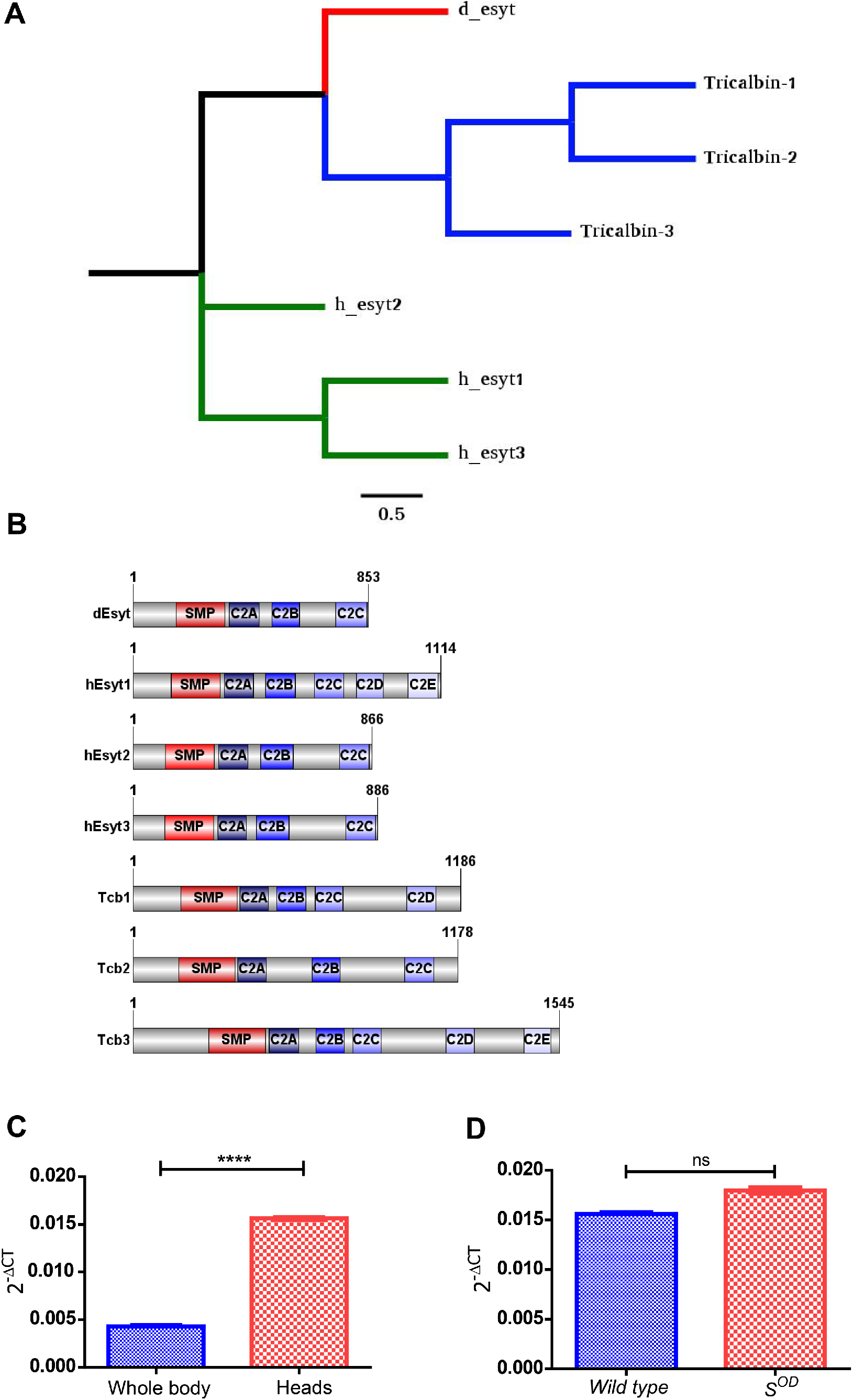
Single gene encoding Esyt expressed in *Drosophila melanogaster.* **(A)** Schematic showing the phylogeny of *Drosophila* Esyt (*dEsyt*), yeast tricalbins (*Tcb1, Tcb2, Tcb3*) and mammalian Esyts (*hEsyt1, hEsyt2,hEsyt3*). **(B)** Schematic of dEsyt-PA protein domain structure aligned with mammalian Esyts and yeast tricalbins (Image generated using IBS, Illustrator for Biological sciences Version 1.0 software http://ibs.biocuckoo.org/) (Liu et al., 2015). Amino acid numbers in each protein are indicated. SMP-SMP domain. C2A, B, C, D, E are the various C2 domains. **(C)** Quantitative real time PCR analysis showing head enrichment of *dEsyt*; the X-axis indicates the wild type tissues from which RNA was isolated and Y-axis indicates the *dEsyt* transcript level expression normalized to the loading control (RP49-Ribosomal protein 49), N=3. **(D)** Quantitative real time PCR analysis showing *dEsy*t transcript level expression in head RNA samples from wild type and *s*^*OD*^. Y-axis indicates the *dEsyt* transcript level expression normalized to the loading control (RP49-Ribosomal protein 49). Bar graphs with mean ± SD are shown. Statistical tests: (C and D) Students unpaired t test (two tailed). ns - Not significant. ****p value <0.0001.

### Generation of *dEsyt^KO^* using CRISPR mediated deletion

To examine a possible function of *dEsyt*, a null allele was generated using CRISPR/Cas9 based genome editing (Kondo and Ueda, 2013). Two sgRNAs were designed in the *dEsyt* genomic region, one sgRNA targeted the 5’ end (approximately 250 bp downstream of the start codon, Target sequence: CCTCTCGGTGGCACGCGACCAGTTGG) and the second sgRNA targeted a region downstream of the second C2 domain in exon 12 (Target sequence: GCTGAAGCAATTCAATATGGAGG). The sites of the sgRNAs were chosen so as to not affect nearby genomic regions implicated in other processes (such as small nuclear RNA U6 gene and gene for tRNA Aspartic acid located near exon 13 of *dEsyt*). The resulting genome edit should remove 488 amino acids of the coding sequence from the 72^nd^ amino acid in the splice variant PA and from the 105^th^ amino acid in the splice variants PB, PC and PD of the *dEsyt* open reading frame which also includes the region encoding the SMP domain and the C2A domain (Fig 2A).

**Figure 2:**
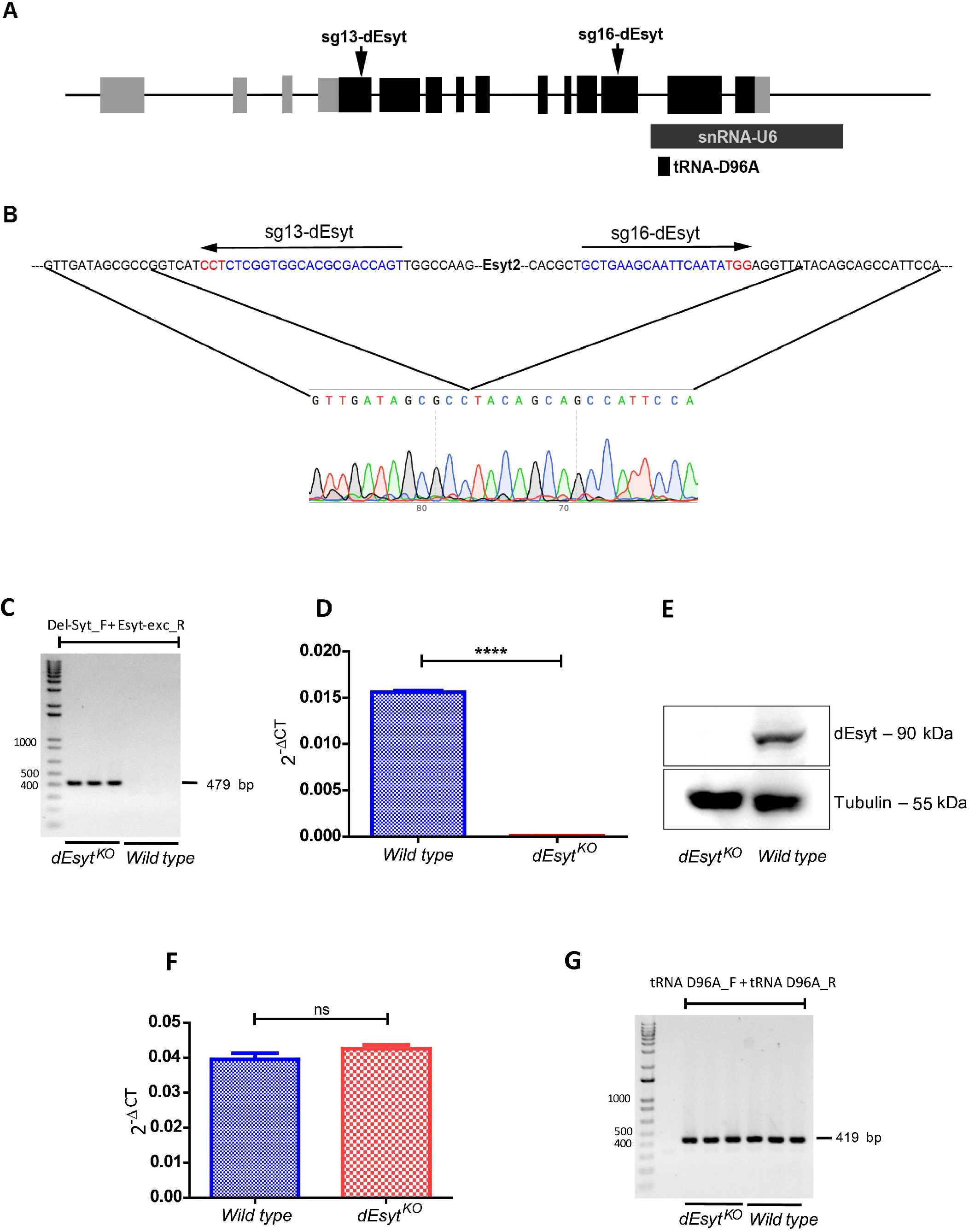
Generation of *dEsyt^KO^* using CRISPR mediated deletion. **(A)** Schematic showing the position of sgRNAs designed to remove 488 aa from different splice variants of the *dEsyt* CDS using CRISPR mediated deletion. Intron/exon structure of the *dEsyt* gene is shown. Coding exons are in black and non-coding exons in grey. Arrows indicate the position of the guide RNAs used in this study for genome engineering. snRNA-U6 and tRNA-D96A indicate the positions of these genes in relation to the *dEsyt* gene. (B) Sanger sequencing chromatogram showing the precise breakpoints induced in the *dEsyt*^*KO*^ allele used in this study. The DNA sequence is shown and the position of the guide RNAs is indicated. **(C)** Image of agarose gel electrophoresis showing the PCR based verification of the knockout. A PCR amplification band of 412 bp is seen in the *dEsyt*^*KO*^ but not in wild type. Primers used for the amplification are shown at the top of the gel and the size of fragments in the DNA ladder is shown in bp. **(D)** Quantitative RT-PCR analysis showing *dEsyt* transcript level expression in the *dEsyt*^*KO*^ and controls. Y-axis indicates the *dEsyt* transcript level expression normalised to the loading control (RP49-Ribosomal protein 49). **(E)** Western blot from head extracts of the genotypes as indicated on the bottom of the blot. The blot was probed with antibody against dEsyt. Tubulin was used as the loading control. **(F)** Comparative real time PCR analysis showing invariable transcript level expression in the small nuclear RNA U6 gene. X-axis shows genotypes and Y-axis indicates the *dEsyt* transcript level expression normalized to the loading control. **(E)** A gel image showing the PCR based verification of the intact genomic region of tRNA D96A in the indicated genotypes. A band of 419 bp is seen in both wild type and *dEsyt*^*KO*^. Bar graphs with mean ± SD are shown. Statistical tests: (C and D) Students unpaired t test (two tailed). ns - Not significant. ****p value <0.0001.

Transgenic flies expressing the dual gRNAs under the U6.2 promoter were crossed to *Act5c-Cas9* virgins. F1 progeny with a deletion of *dEsyt* gene was identified using a PCR based strategy obtaining DNA from heterozygotes at the F1 generation. Seven F1 progeny tested positive for this deletion; two with the brightest band (indicative of maximum number of cells with deletion) were further crossed to third chromosome balancers. 30 F2 progeny were screened for deletion and four of the pure lines were maintained as final heterozygous stocks, after confirming the absence of the dual gRNA and Cas9 transgenes. The genomic boundaries of the deletion were estimated by PCR based amplification with primers flanking the *dEsyt* genomic region. The mutant allele generated a band of 412 bp as against 3.7 kb region in controls (Fig 2C). Sanger sequencing of this PCR product was used to confirm the exact breakpoints (Fig 2B); this allele is referred to as *dEsyt*^*KO*^.

*dEsyt*^*KO*^ was validated using RT-qPCR on total head RNA from homozygous flies- this showed negligible amounts of *dEsyt* transcripts in *dEsyt*^*KO*^ compared to controls (Fig 2D). Using an antibody against dEsyt for immunoblot analysis (Kikuma et.al 2017), *dEsyt*^*KO*^ was confirmed to be a protein null allele (Fig 2E). No change was detected in the transcript levels of the small nuclear RNA U6 gene (Fig 2F) and the genomic region of tRNA D96A (Fig 2G) in *dEsyt*^*KO*^.

### *dEsyt* is dispensable for normal phototransduction

During larval development, *dEsyt*^*KO*^ homozygous larvae show a delay in development and adult flies that eclose are smaller than controls (Sup Fig S1A, B). These phenotypes can be rescued by pan larval reconstitution with *dEsyt* cDNA (Sup Fig S1C, D). Although most *dEsyt*^*KO*^ homozygotes do not eclose as adults, when grown on nutrient enriched medium, a small proportion is viable as homozygous adults.

To test the potential role of *dEsyt* in supporting lipid transfer at MCS, we studied the phenotypes of these homozygous *dEsyt*^*KO*^ in adult photoreceptors. We measured electrical responses to light in adult flies. Electroretinograms (ERG) were recorded from wild type and *dEsyt*^*KO*^ adults of matched age and eye colour. We found that the absolute ERG response of *dEsyt*^*KO*^ was smaller than that of controls (Fig 3A). However the ERG response is an extracellular recording that scales with the size of the eye. Since *dEsyt*^*KO*^ are smaller in size than controls, we normalized the raw ERG response of each animal to its body size (Fig 3B); this normalized ERG amplitude was not significantly different between control and *dEsyt*^*KO*^.

**Figure 3:**
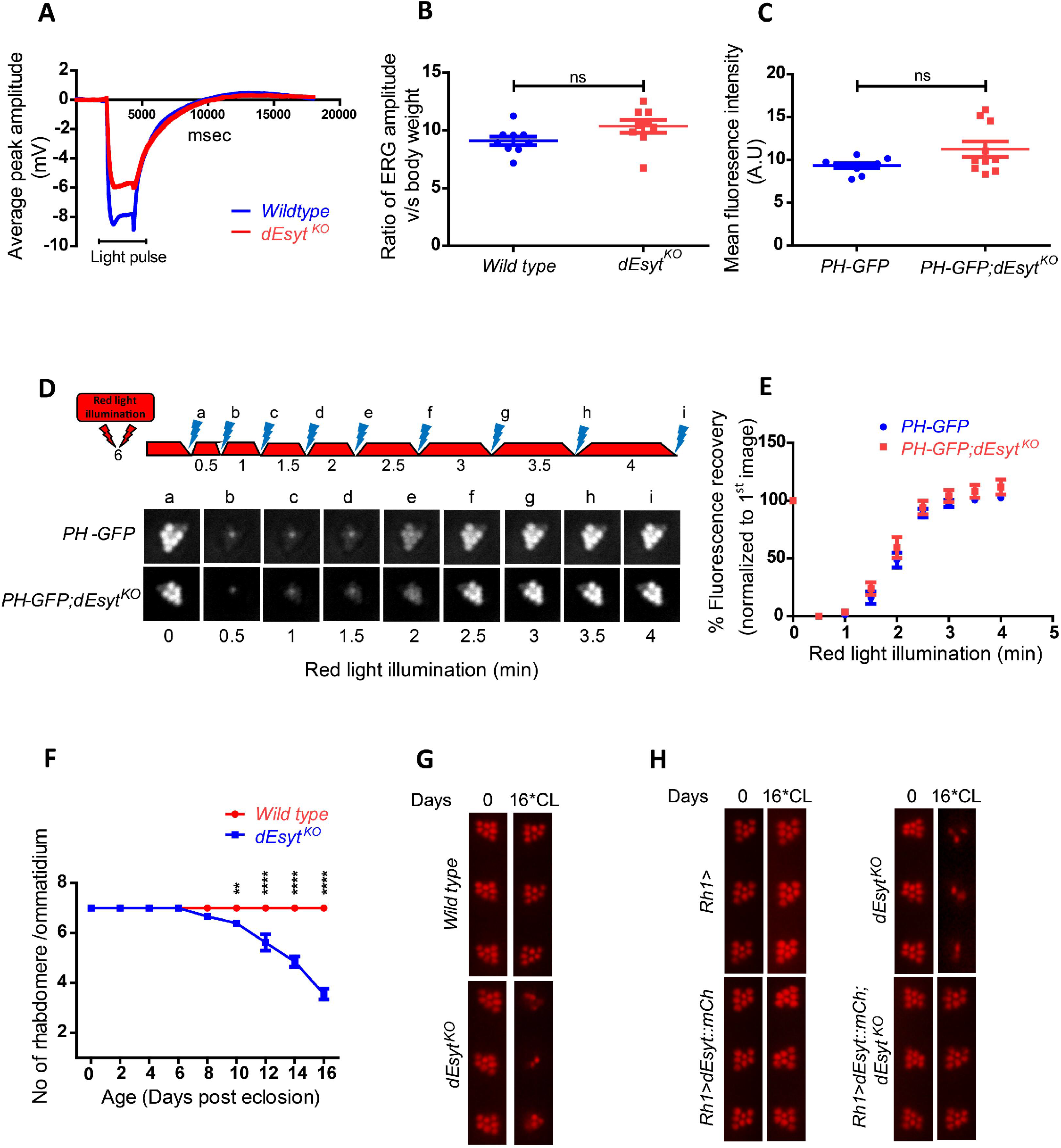
*dEsyt* is dispensable for phototransduction. **(A)** Representative ERG trace from dark reared 0-1 day old flies of the indicated genotype stimulated by 2s flash of green light. The duration of the stimulating light is shown. Y-axis shows the amplitude of the ERG response. X-axis shows the duration of the recording. **(B)** Graph showing ERG amplitude normalized to body weight. Y-axis shows the ratio of ERG amplitude of individual flies to their body weight. X-axis indicates genotypes. Each data point represents an individual fly tested. Error bar represents s.e.m. **(C)** Quantification of the mean fluorescence intensity of the deep pseudopupil formed by the PIP_2_ probe PH-GFP in the one day old flies of the indicated genotypes (n=10). **(D)** Deep pseudopupil imaging of PIP_2_ levels in the microvillar membrane of photoreceptors. The fluorescence of the PH-GFP probe is depicted. The protocol used is shown with red light illumination periods shown as red bars and flashes of blue light (a-f) used for image capture depicted. Representative deep pseudopupil images acquired at specified time points are depicted. Genotypes as indicated. **(E)** Graph quantifying the recovery kinetics of the fluorescent pseudopupil with time, X-axis represents the genotypes, Y-axis represent intensity normalised to the intensity of the first image. (n=10). Error bars represent s.e.m. **(F)** Quantification of the time course of retinal degeneration. 10 ommatidia from 5 separate flies of each genotype were scored and plotted. Y-axis is the number of intact rhabdomeres/ommatidium. The maximum value possible is 7. X-axis is the age of the flies post eclosion. **(G)** Representative optical neutralization images showing rhabdomere integrity of the indicated genotypes. Rearing conditions and the age of the flies are indicated on top. (*CL-Constant Light) **(H)** Representative ON images showing the rescue of rhabdomere structural integrity in *dEsyt*^*KO*^ mutants on expressing the dEsyt∷mCherry transgene. Genotypes of the flies are indicated on the left. Rearing conditions and the age of the flies are indicated on top. Scatter plots and XY plots with mean ± SD are shown. Statistical tests: (B and C) Student’s unpaired t test. (E and F) Two-Way ANOVA Grouped analysis with Bonferroni post-tests to compare replicate means. ns - Not significant; **p value < 0.01; ****p value <0.0001.

Since contact sites are proposed to support lipid transfer reactions during cell signaling, we tested the requirement for dEsyt in supporting PIP_2_ turnover during phototransduction. We compared the level of PIP_2_ at the microvillar PM of photoreceptors in wild type and *dEsyt*^*KO*^ flies. To monitor the changes in the PIP_2_ levels, the PH domain of PLCδ fused to GFP (hereafter referred to as PH-GFP) was utilized (Yadav et al., 2015). This analysis revealed no difference in the basal levels of PIP_2_ between controls and *dEsyt*^*KO*^ (Fig 3C). Further, following bright light stimulation to activate PLCβ and deplete PIP_2_ at the microvillar PM, we found no difference in the rate at which PIP_2_ levels were restored at the microvillar PM (Fig 3D, E) in *dEsyt*^*KO*^. Similarly, the P4M-GFP probe was utilised to monitor the levels of PI4P which acts as the substrate for *dPIP5K* to produce PIP_2_ (Balakrishnan et al., 2018); basal PI4P levels and PI4P turnover kinetics were similar between wild type and *dEsyt*^*KO*^ (Sup Fig S2A, B, E).

Light dependent retinal degeneration is a common outcome when genes essential for phototransduction are disrupted (Raghu et al., 2012). To test the role if any, of *dEsyt* in regulating phototransduction, we monitored the integrity of rhabdomere (apical PM) structure following illumination. Although normal at eclosion, when flies were grown under constant bright illumination, normal photoreceptor ultrastructure was retained until 10 days post-eclosion. However, photoreceptors of *dEsyt*^*KO*^ underwent progressive retinal degeneration when observed till day 16 with substantial loss of rhabdomere integrity (Fig 3F, G). This light dependent degeneration exhibited by the mutant *dEsyt* homozygotes was rescued by reconstitution with wild type dEsyt transgene (Fig 3H).

### Loss of dEsyt enhances *rdgB^9^* phenotypes

It is reported that mammalian Esyt1 enhances the activity of Nir2 by recruiting it to the ER-PM junctions subsequent to an elevation in cytosolic calcium levels (Chang et al., 2013). We tested the function of dEsyt in supporting the activity of RDGB, the *Drosophila* ortholog of Nir2. For this, we generated a double mutant strain of *dEsyt*^*KO*^ with *rdgB*^*9*^, a hypomorphic allele of *rdgB* (*rdgB*^*9*^*;dEsyt*^*KO*^). Using this strain, we compared the impact of *dEsyt* deletion on the phenotypes of the *rdgB*^*9*^ single mutant.

In *Drosophila*, *rdgB* mutants show a number of key phenotypes including light-dependent retinal degeneration, reduced electrical response to light and altered PIP_2_ homeostasis at the microvillar PM (Yadav, et.al 2015). We compared the time course of retinal degeneration in *rdgB*^*9*^ v *rdgB*^*9*^*;dEsyt*^*KO*^ and noted that *dEsyt*^*KO*^ accelerates the time course of retinal degeneration in *rdgB*^*9*^ (Fig 4A i, ii and 4B). Over this time period, *dEsyt*^*KO*^ itself does not show any retinal degeneration; the retinal degeneration of the *dEsyt*^*KO*^ occurs much later after eclosion.

**Figure 4:**
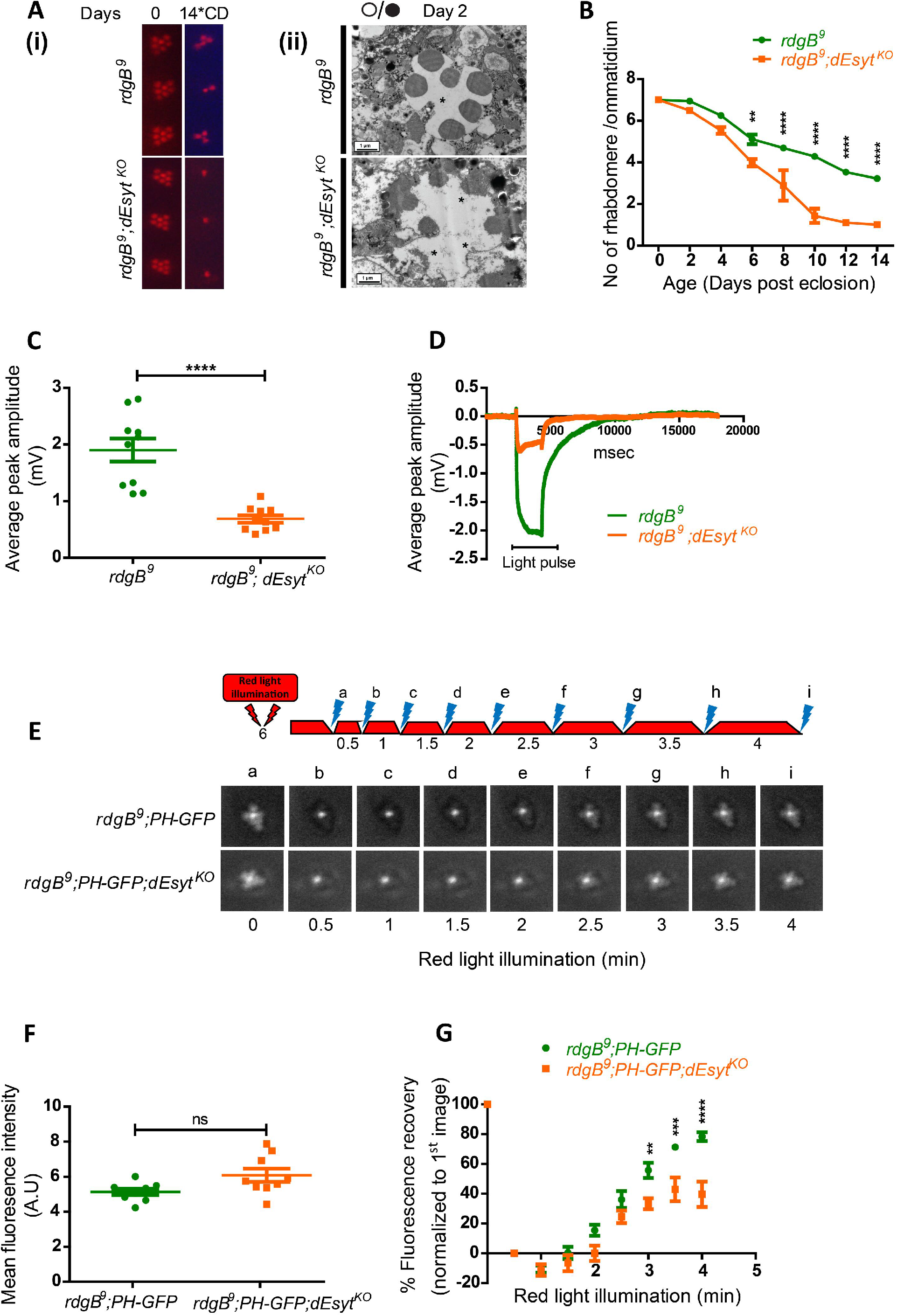
Loss of *dEsyt* enhances *rdgB* mutant phenotypes. **(A) (i)** Representative optical neutralization (ON) images showing rhabdomere structure of the indicated genotypes. Rearing conditions and the age of the flies are indicated on top. (*CD- Constant Dark). (ii) TEM images showing a single ommatidium from the photoreceptors of *rdgB*^*9*^ and *rdgB*^*9*^*;dEsyt*^*KO*^ mutants reared in 12 hr L/D cycle (⚫/⚪) for 2 days. Scale bar: 1 μM. (*- degenerated rhabdomere) **(B)** Quantification of the time course taken for retinal degeneration. 10 ommatidia were scored from 5 flies of each genotype and plotted. **(C)** Representative ERG traces showing the duration of light pulse, X-axis indicates time in msec and Y-axis indicates the average ERG amplitude in mV. **(D)** Graph showing ERG amplitude of *rdgB*^*9*^ and *rdgB*^*9*^*;dEsyt*^*KO*^ normalized to body weight. Y-axis shows the ratio of ERG amplitude of individual flies to their body weight. X-axis indicates genotypes. Each data point represents an individual fly tested. Error bar represents s.e.m. **(E)** Deep pseudopupil imaging of PIP_2_ levels in the microvillar membrane of photoreceptors. The fluorescence of the PH-GFP probe is depicted. The protocol used is shown with red light illumination periods shown as red bars and flashes of blue light (a-f) used for image capture depicted. Representative deep pseudopupil images acquired at specified time points are depicted. Genotypes as indicated. **(F)** Quantification of the mean fluorescence intensity of the PIP_2_ probe PH-GFP from the deep pseudopupil formed by one day old flies of the indicated genotypes (n=10). **(G)** Graph quantifying the recovery kinetics of the fluorescent pseudopupil with time, X-axis represents the genotypes, Y-axis represent intensity normalized to the intensity of the first image. (n=10). Scatter plots and XY plots with mean ± SD are shown. Statistical tests: (C and F) Student’s unpaired t test. (B and G) Two-Way ANOVA Grouped analysis with Bonferroni post-tests to compare replicate means. ns - Not significant; **p value < 0.01; ***p value < 0.001; ****p value <0.0001.

We also studied the ERG amplitude in response to a 1s flash of light. As previously reported (Yadav et al., 2015), *rdgB*^*9*^ mutants show a reduced ERG amplitude and this was further reduced in *rdgB*^*9*^*;dEsyt*^*KO*^ (Fig 4C, D). Finally we also measured PIP_2_ levels at the microvillar PM under resting conditions. The PIP_2_ levels in *rdgB*^*9*^ are lower than in wild type (Yadav et.al, 2015) and there was no difference in resting PIP_2_ levels between *rdgB*^*9*^ and *rdgB*^*9*^*;dEsyt*^*KO*^ (Fig 4F). However, we also measured the rate of recovery of microvillar PIP_2_ following its depletion by strong PLCβ activation. Under these conditions, the reduced rate of PIP_2_ recovery in *rdgB*^*9*^ was further delayed in *rdgB*^*9*^*;dEsyt*^*KO*^ (Fig 4E, G) and a similar observation was also made for PI4P levels (Sup Fig S2C, D, E). Collectively, loss of *dEsyt* worsened the three key functional and biochemical outcomes in a hypomorphic allele of *rdgB*.

### *dEsyt* is localized to ER-PM contact sites in photoreceptors

To understand the mechanism by which loss of dEsyt enhances *rdgB*^*9*^ phenotypes, it was crucial to decipher the localization of dEsyt. Since Esyt proteins have been described as components of MCS in both yeast and mammalian cells, we tested if the same is also true in *Drosophila* photoreceptors. For this we labelled wild type retinae with anti-dEsyt serum (Kikuma et al., 2017). Single confocal images of photoreceptor cross-sections (R1-R6) revealed a crescent shaped distribution of dEsyt at the MCS between the microvillar PM and the SMC (Fig 5B); this staining was absent in *dEsyt*^*KO*^ photoreceptors demonstrating the specificity of the labelling detected. To further confirm this observation, we expressed dEsyt tagged to the fluorescent protein mCherry in photoreceptors and examined its localization using an antibody against mCherry; here too we found that dEsyt∷mCherry was localized to the MCS at the base of the microvilli in photoreceptors (Fig 5A); staining was not seen in wild type photoreceptors not expressing dEsyt∷mCherry indicating the specificity of the staining. Collectively, these findings demonstrate that dEsyt is localized at the ER-PM MCS in photoreceptors.

**Figure 5:**
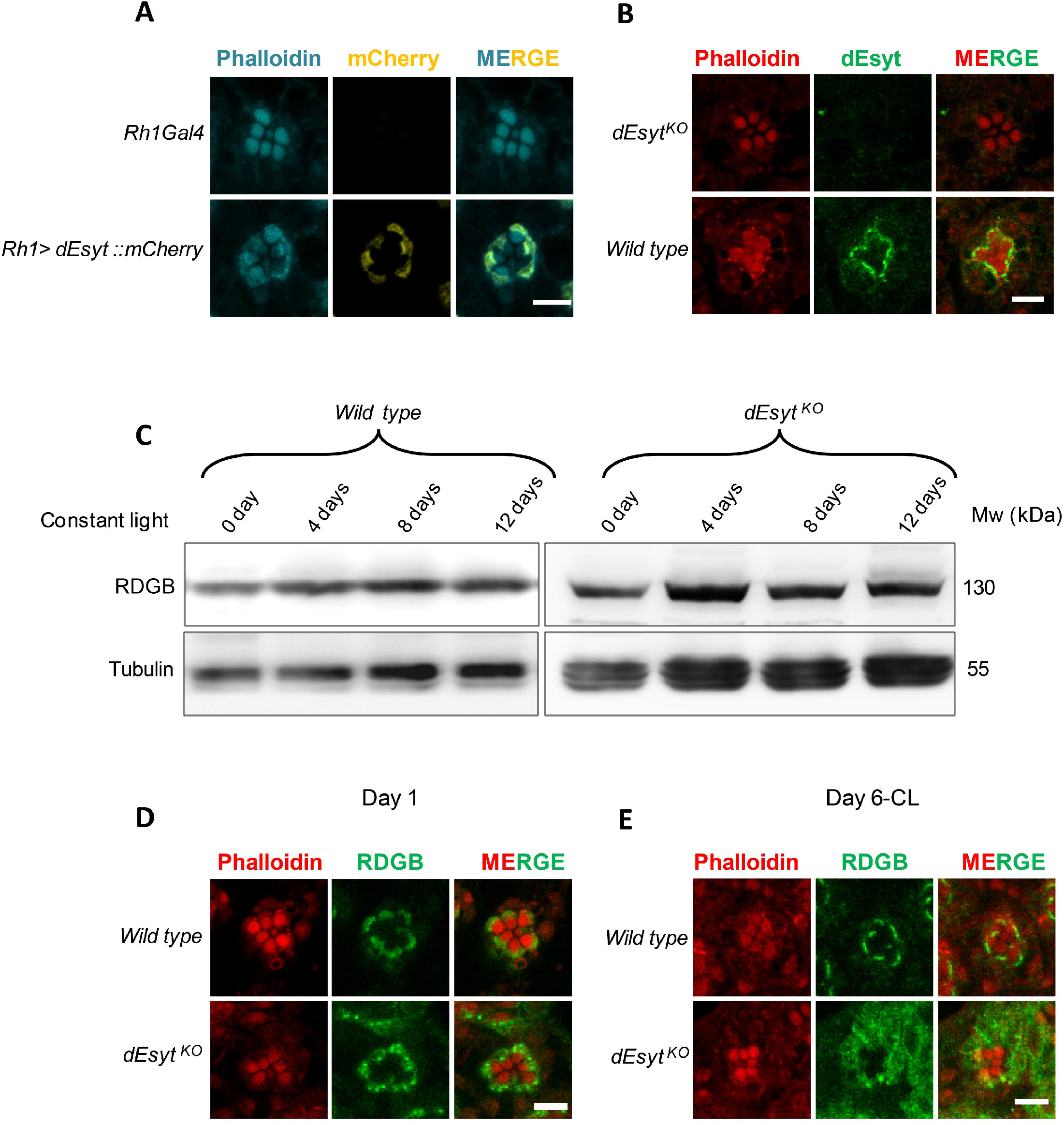
dEsyt localizes to MCS and determines RDGB localization. **(A)** Confocal images showing the localization of exogenously expressed dEsyt∷mCherry protein expressed using the eye specific Rh1-Gal4 in one day old dark reared flies. Rh1-Gal4 is shown as a control. A single ommatidium is shown. Scale bar: 5 μM. Phalloidin marks F-actin staining and highlights rhabdomeres R1-R7. **(B)** Confocal images showing the localization of endogenous dEsyt protein in photoreceptors of one day old dark reared flies probed with antibody against dEsyt. Scale bar: 5 μM. *dEsyt*^*KO*^ shows no staining when probed with dEsyt antibody. **(C)** Western blot from the head extracts of wild type and *dEsyt*^*KO*^ probed with the antibody against RDGB. Rearing conditions, age of the flies and genotype is indicated on top of the blot. Tubulin was used as the loading control. Confocal images showing the localization of RDGB in wild type and *dEsyt*^*KO*^ photoreceptors of flies which are **(D)** 1 day old-dark reared **(E)** 6 days old-exposed to constant illumination. For (D) and (E) RDGB visualized is detected using an antibody against the endogenous protein. Rhabdomeres are outlined using Phalloidin which marks F-actin.

### dEsyt function is required to localize RDGB to ER-PM contact sites

One mechanism by which loss of *dEsyt* might enhance *rdgB*^9^ phenotypes is by reducing the levels of the endogenous RDGB protein. We checked the levels of RDGB protein in *dEsyt*^*KO*^ mutants exposed to bright light illumination for increasing time intervals and found that the RDGB protein levels remain unaltered even after retinal degeneration is initiated in *dEsyt*^*KO*^ (Fig 5C). A second mechanism could be the mislocalization of the RDGB protein away from the ER-PM MCS. In wild type *Drosophila* photoreceptors, RDGB is strictly localized to the MCS and we found that this localization was not disrupted in newly eclosed *dEsyt*^*KO*^ flies (Fig 5D). However, when exposed to constant illumination for 6 days post eclosion, *dEsyt*^*KO*^ photoreceptors showed mislocalization of RDGB away from the MCS; the protein was now diffused across the photoreceptor cell body (Fig 5E). By contrast, under the same conditions, RDGB localization to the MCS was retained in wild type photoreceptors (Fig 5E). Thus dEsyt function is required for the normal localization of RDGB at the MCS in photoreceptors.

### dEsyt is necessary for PLCβ-dependent modulation of ER-PM MCS

The most direct way of assessing MCS structure is by transmission electron microscopy (TEM). We used fixation methods that allowed us to enhance membrane preservation and were able to visualize MCS clearly in photoreceptors (Fig 6A i, ii). Such images allowed us to quantify the fraction of PM at the base of the microvilli that was in contact with the SMC (hereafter referred to as MCS density) in each ultrathin section of a photoreceptor. On day 1 after eclosion, in R1-R6 photoreceptors of dark-reared wild type flies, the MCS density was ca. 50% (Fig 6A i, ii and 6B i). As flies aged, the MCS density decreased to ca. 30% by day 14 (Fig 6A iii, iv and 6B i). Under the same conditions, the MCS density in the UV sensitive R7 photoreceptor remained constant as a function of age (Fig 6B ii). We also measured the MCS density in the *norpA*^*P24*^ mutant allele that lacks PLCβ protein expression (Pearn et al., 1996). MCS density in *norpA*^*P24*^ was ca. 30% at eclosion (Fig 6C i, ii) but in contrast to wild type flies, *norpA*^*P24*^ flies did not show any reduction as a function of age; i.e day 14 *norpA*^*P24*^ photoreceptors showed the same MCS density as wild type cells at eclosion (Fig 6D i). Thus MCS density in photoreceptors is dependent on age and PLCβ activity.

**Figure 6:**
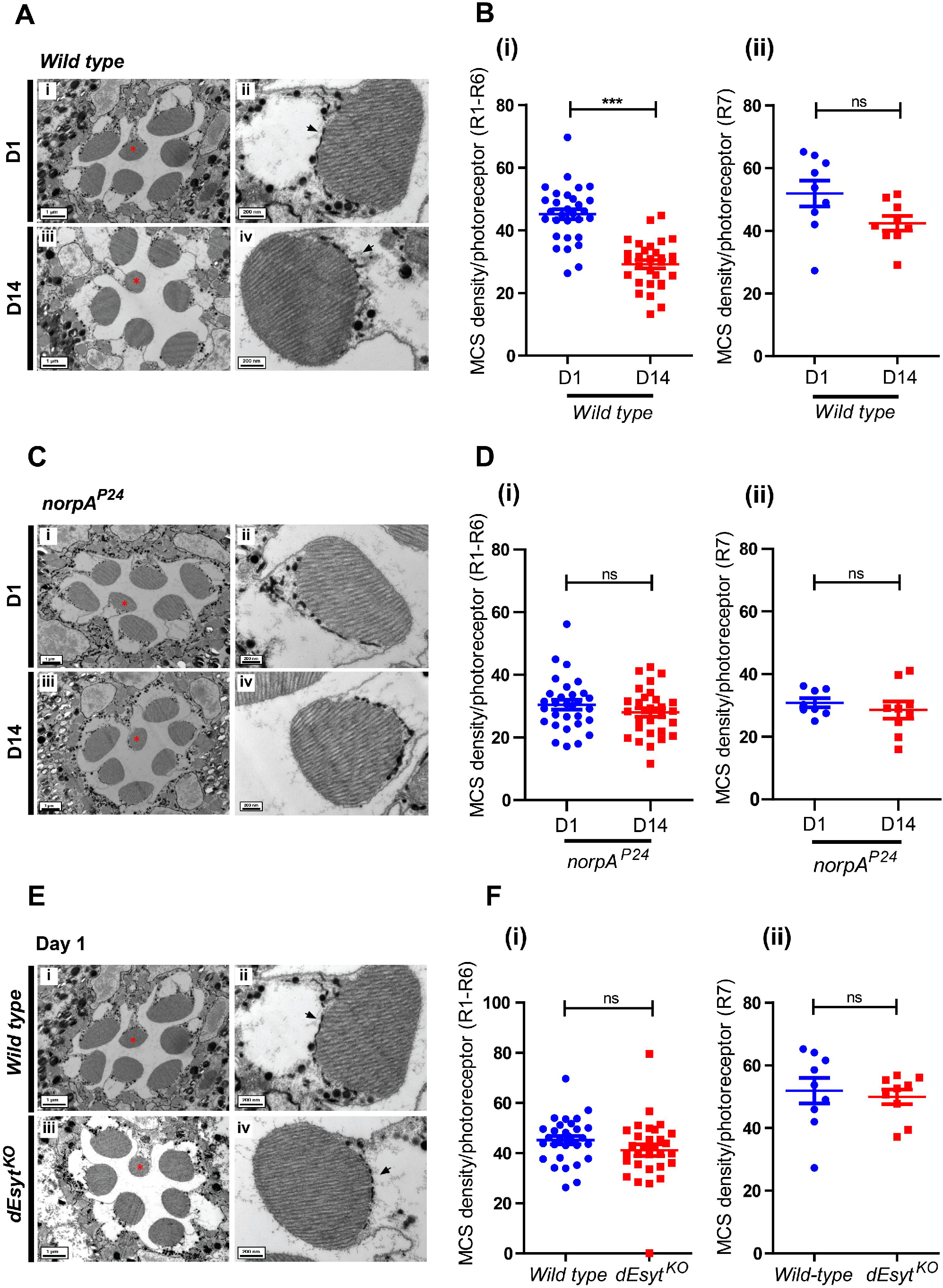
dEsyt stabilizes MCS in photoreceptors. **(A)** TEM images of a single ommatidium from wild type photoreceptors of flies reared in dark **(i)** day 1 (D1) and **(iii)** day 14 (D14), scale bar: 1 μM. (* - R7 photoreceptor). Magnified image showing a single photoreceptor from the same ommatidium **(ii)** day 1 **(iv)** day 14. Scale bar: 200 nm. Arrow indicates the SMC forming an MCS with the microvillar PM. **(B)** Quantification of the number of MCS per wild type photoreceptor of day 1 and day 14 old flies reared in dark (from figure 6A), Y-axis indicates the number of MCS/photoreceptor. n=30 photoreceptors from 3 separate flies (R1-R6) for **(i)**. n=9 photoreceptors from 3 separate flies (R7) in **(ii).** **(C)** TEM images of a single ommatidium from *norpA*^*P24*^ photoreceptors of flies reared in dark **(i)** day 1 and **(iii)** day 14, scale bar: 1μM. (* - R7 photoreceptor). Magnified image showing a single photoreceptor from the same ommatidium **(ii)** day 1 **(iv)** day 14. Scale bar: 200 nm. **(D)** Quantification of the number of MCSs per photoreceptor (from figure 6C), X-axis indicates the genotype and age of the flies and Y-axis indicates the number of MCS/photoreceptor **(i)** n=30 photoreceptors from 3 separate flies (R1-R6) **(ii)** n=9 photoreceptors from 3 separate flies (R7); **(E)** TEM images of a single ommatidium from **(i)** wild type and **(iii)** *dEsyt*^*KO*^ photoreceptors of 0-1 day old flies reared in dark, scale bar: 1 μM. (* - R7 photoreceptor) **(ii, iv)** Magnified image showing a single photoreceptor from the ommatidium image shown on the left. Scale bar: 200 nm. **(F)** Quantification of the number of MCS per photoreceptor of wild type day 1 v/s *dEsyt*^*KO*^ day 1 old flies reared in dark (from figure 6C), X-axis indicates the genotype and age of the flies and Y-axis indicates the number of MCS/photoreceptor **(i)** n=30 photoreceptors from 3 separate flies (R1-R6) **(ii)** n=9 photoreceptors from 3 separate flies (R7). Scatter plots with mean ± SD are shown. Statistical tests: (B, D and F) Student’s unpaired t test. ns - Not significant; ***p value <0.001.

To assess the contribution of dEsyt to ER-PM MCS structure, we performed TEM on *dEsyt*^*KO*^ photoreceptors to visualize and quantify MCS density (Fig 6E iii, iv). At eclosion, MCS density was not different between wild type and *dEsyt*^*KO*^ for R1-R6 photoreceptors (Fig 6E i-iv and 6F i); a similar result was seen in UV sensitive R7 photoreceptors (Fig 6F ii). Thus loss of dEsyt does not impact the establishment of ER-PM contact sites during development in *Drosophila* photoreceptors. We then examined MCS density as a function of age in *dEsyt*^*KO*^ (Fig7Ai-iv). While MCS density was reduced from ca. 50% (day 1) to 30% at day 14 in wild type flies (Fig 6B i), in *dEsyt*^*KO*^ flies, the number of MCS was reduced to almost none by day 14 (Fig 7B i). By contrast, in UV sensitive R7 photoreceptors, MCS density was only modestly reduced in *dEsyt*^*KO*^ (Fig 7B ii).

**Figure 7:**
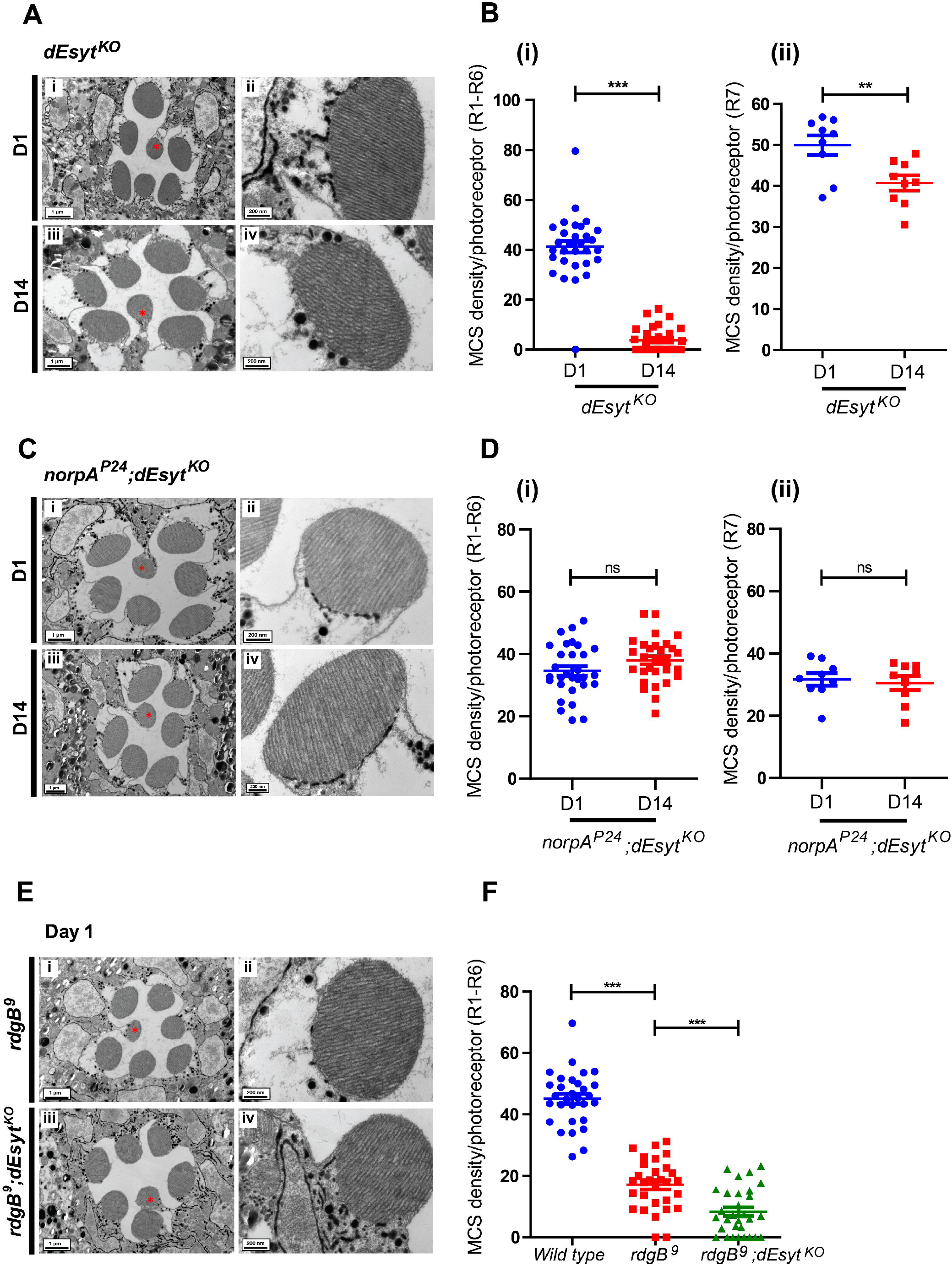
dEsyt is necessary for PLCβ-dependent modulation of ER-PM MCS. TEM images of a single ommatidium from photoreceptors of flies reared in dark, scale bar: 1 μM. (* - R7 photoreceptor). **(A) (i)** *dEsyt*^*KO*^- day 1 **(iii)** *dEsyt*^*KO*^- day 14 **(C) (i)** *norpA*^*P24*^*;dEsyt*^*KO*^- day 1 **(iii)** *norpA*^*P24*^*;dEsyt*^*KO*^- day 14 **(E) (i)** *rdgB*^*9*^ - day 1 and **(iii)** *rdgB*^*9*^*;dEsyt*^*KO*^ - day 1 **(A-ii, iv; C-ii, iv; E-ii, iv)** Magnified image showing a single photoreceptor from the ommatidium image shown on the left. Scale bar: 200 nm. **(B, D)** Quantification of the number of MCS per photoreceptor, X-axis indicates the genotype and age of the flies and Y-axis indicates the number of MCS/photoreceptor **(i)** n=30 photoreceptors from 3 separate flies for R1-R6. **(B & D ii)** n=9 photoreceptors from 3 separate flies for R7. **(F)** Quantification of the number of MCS per photoreceptor for R1-R6 in 1 day old dark reared wild type (from Figure 6B i), *rdgB*^*9*^ and *rdgB*^*9*^*;dEsyt*^*KO*^ flies. n=30 photoreceptors from 3 separate flies. Scatter plots with mean ± SD are shown. Statistical tests: (B and D) Student’s unpaired t test. (F) one-way ANOVA with post hoc Tukey’s multiple pairwise comparison. Ns -not significant, **p value <0.01; ***p value <0.001.

We tested the impact of PLCβ activity on the enhanced loss of MCS in *dEsyt*^*KO*^. For this, we compared MCS density in *dEsyt*^*KO*^ with that in the *norpA*^*P24*^*;dEsyt*^*KO*^ double mutant (Fig 7C i-iv). Strikingly, the absence of basal PLCβ activity in *norpA*^*P24*^*;dEsyt*^*KO*^ photoreceptors completely rescued MCS number in 14 day flies; this was in sharp contrast to *dEsyt*^*KO*^ flies which at this stage typically shows very few or no MCS (Fig 7C iii, iv and 7D i). Thus the loss of MCS in *dEsyt*^*KO*^ with age depends on ongoing PLCβ activity.

We also examined MCS structure in *rdgB*^*9*^ photoreceptors (Fig 7E i, ii). In dark reared flies, just after eclosion, MCS density in R1-R6 photoreceptors was already reduced significantly (Fig 7F i). *rdgB*^*9*^*;dEsyt*^*KO*^ photoreceptors also showed very few MCS in the photoreceptors (Fig 7E iii, iv); these were quantified only from photoreceptors in which there was minimal degeneration of rhabdomeres (Fig 7E iii). Further, *rdgB*^*9*^*;dEsyt*^*KO*^ double mutants showed substantially greater rhabdomeral degeneration than *rdgB*^*9*^ (Fig 4A ii) when reared for 2 days under conditions of 12 hr L/D cycle reflecting the ultrastructural level observations seen in the optical neutralization assays (Fig 4B).

## Discussion

Studies in yeast and mammalian cells have identified proteins that when overexpressed, localize to ER-PM MCS. However, in most cases, surprisingly, loss-of-function of these proteins appear to not impact cellular function or *in vivo* physiology. Though Esyt can localize to ER-PM MCS in cultured mammalian cells (Chang et al., 2013; Giordano et al., 2013), depletion of Esyt in cultured mammalian cells apparently does not impact cell physiology and knockout of Esyt in mice does not have discernible impact on animal physiology (Sclip et al., 2016; Tremblay and Moss, 2016) raising questions on their role *in vivo*. However, in this study, we found that a null mutant of the only Esyt in *Drosophila* showed developmental delay, reduced adult viability and homozygotes that eclosed were smaller than controls; the reduced body size could be rescued by reconstitution with a wild type transgene suggesting an important role for *dEsyt* in growth. While the cellular and molecular basis of these phenotypes remains to be established, these data clearly demonstrate a function for Esyt in supporting animal physiology *in vivo*.

Importantly, we analysed photoreceptor structure and function in *dEsyt*^*KO*^ since this cell type critically depends on lipid transfer activity at its ER-PM MCS for phototransduction (Yadav et al., 2016, 2018). In newly eclosed *dEsyt*^*KO*^ flies, photoreceptor ultrastructure was unaffected and ER-PM MCS density was not different from that in wild type flies. Thus dEsyt is not required during development to establish ER-PM MCS. We also found that *dEsyt*^*KO*^ showed normal electrical responses to light and basal levels of PM PI4P and PIP_2_ as well as their turnover during PLCβ signaling was unaffected; this finding implies that dEsyt function is dispensable for PIP_2_ turnover (that in turn depends on PI transfer at ER-PM MCS) during PLCβ signaling. Our observation is consistent with that reported for mammalian Esyt that the SMP domain does not exhibit specific PI transfer activity *in vitro* (Saheki et al., 2016; Schauder et al., 2014), that loss of Esyt function in cultured mammalian cells does not impact PIP_2_ turnover during PLCβ signaling and that depletion of Esyt does not impact store-operated calcium influx, a key output of PLCβ activation (Giordano et al., 2013).

Despite the apparent lack of a phenotype in *dEsyt*^*KO*^ photoreceptors, we found that loss of dEsyt enhances all three key phenotypes of *rdgB*^*9*^, a hypomorphic allele with ca 5% residual RDGB protein in photoreceptors. Importantly, the impact of *dEsyt*^*KO*^ on the reduced light response and PM PIP_2_ levels in *rdgB*^*9*^ occurred in newly eclosed flies, prior to the onset of retinal degeneration. Conceptually, this enhancement could be due to a change in the levels of the major phototransduction proteins but this was ruled out since we found no alteration in the levels of these proteins (Rhodopsin, PLC, Gq and TRP) in *rdgB*^*9*^*;dEsyt*^*KO*^ (Sup Fig S3). Interestingly, we found that *dEsyt*^*KO*^ did not enhance the reduced ERG amplitude resulting from the depletion of dPI_4_KIIIα (Sup Fig S4), the enzyme implicated in the conversion of PI to PI4P at the PM of photoreceptors (Balakrishnan et al., 2018). Thus the enhancement of *rdgB*^*9*^ phenotypes is specific and not seen even in cells depleted of the immediate next component of the PIP_2_ cycle in photoreceptors. These findings strongly indicate a key role for dEsyt in specifically supporting the function of RDGB at the ER-PM MCS. Our data suggest that in photoreceptors with a full capacity of RDGB function, cells can dispense with the requirement for dEsyt. However in cells with limiting RDGB function (such as *rdgB*^*9*^), the requirement for dEsyt function becomes apparent. Thus our data clearly support a role for dEsyt in supporting the lipid transfer function of RDGB at ER-PM contact sites *in vivo*. Consistent with this model, we found that the dEsyt protein is localized to the base of the microvillar PM at the ER-PM contact site where the RDGB protein is also specifically localized (Vihtelic et al., 1993; Yadav et al., 2018).

Although we found a normal density of ER-PM MCS in *dEsyt*^*KO*^ photoreceptors at eclosion, we noted that this changed as a function of age and illumination. Wild type photoreceptors show a reduction in ER-PM MCS density with age; however, the drop in MCS density under the same conditions was much greater in *dEsyt*^*KO*^ with virtually no MCS observed in 14 day old, constant dark reared flies. These findings imply that dEsyt function is essential to maintain the stability of ER-PM MCS in photoreceptors. We also observed that although the RDGB protein was correctly localized at the MCS at eclosion, in *dEsyt*^*KO*^ photoreceptors, the protein became delocalized as a function of age in the absence of dEsyt function. By day 6 the RDGB protein was substantially delocalized from the base of the microvilli in *dEsyt*^*KO*^, although the total amount of RDGB protein in the cell was not reduced. During this study, we also noted that the MCS density was lower in *rdgB*^*9*^*;dEsyt*^*KO*^ compared to *rdgB*^*9*^ alone. This observation offers an explanation for the ability of *dEsyt*^*KO*^ to enhance the physiological phenotypes of *rdgB*^*9*^ on day 1 post-eclosion; i.e the reduced MCS density in *rdgB*^*9*^ with limited lipid transfer activity is further compromised by the additional loss of MCS resulting from *dEsyt*^*KO*^. Taken together, these findings suggest that in the absence of dEsyt function, it is likely that RDGB (and perhaps other MCS localized proteins) become delocalized leading to disruption of its lipid transfer function. These findings lead us to a model of dEsyt function in photoreceptors where its key role is to stabilize ER-PM MCS and facilitate the correct localization of relevant proteins at this MCS.

Despite intense interest in the function of MCS, very little is known about the mechanisms by which MCS density is tuned in response to signals. In photoreceptors, a key signal is light and this is transduced through G-protein coupled PLCβ activity [reviewed in (Raghu et al., 2012)]. In this study, we made several observations that MCS density is linked to PLCβ activity: (i) in newly eclosed flies MCS density was reduced in *norpA* mutants that lack PLCβ activity (ii) in wild type photoreceptors, ER-PM MCS density decreased as a function of age and this decrease was dependent on ongoing PLCβ activity. (iii) the loss of MCS in *dEsyt*^*KO*^ photoreceptors could be suppressed by *norpA* mutants. Together, these findings suggest that PLCβ activity is a key element of the mechanism by which MCS density is regulated. Interestingly, in *rdgB*^*9*^, in which PM PIP_2_ levels are lower than wild type, MCS density was reduced. Since both PLCβ and RDGB regulate PIP_2_ levels, our observations suggest that MCS density in photoreceptors is modulated by PIP_2_ turnover at the PM. Since PIP_2_ turnover is a key step of phototransduction in *Drosophila* photoreceptors, it is possible that in this cell type, illumination modulates MCS density. In turn this will alter the biochemical activity of proteins localized at the MCS and their ability to maintain microvillar PM homeostasis. Future studies will likely reveal the molecular mechanism by which MCS density is tuned to ambient illumination and the broader question of PM homeostasis by MCS in eukaryotic cells.

## Supporting information

Esyt_Sup fig

## Acknowledgements

This work was supported by the National Centre for Biological Sciences-TIFR, a Wellcome-DBT India Alliance Senior Fellowship (IA/S/14/2/501540) to PR, a Wellcome-DBT India Alliance Early Career Fellowship (IA/E/17/1/503653) to SM and the Department of Biotechnology, Government of India (BT/PRJ3748/GET/l 19/27/2015). We thank the *Drosophila* Facility, Central Imaging Facility and Electron Microscopy Facility at NCBS for support. We thank Dr. D. Dickman for sharing valuable reagents.

## Materials and methods

### Fly stocks

Flies (*Drosophila melanogaster*) were raised on rich medium (composition : corn flour, black jaggery, agar, dry yeast, propionic acid, methyl parahydroxy benzoate, orthophosphoric acid) at 25°C maintaining 50% relative humidity in a constant temperature laboratory incubator. The environment within the incubator was completely devoid of illumination except for the brief pulses of light to which the flies were subjected when the incubator doors were opened. The experiments were carried out in three different conditions: constant illumination, constant dark and 12 hour light-dark cycle. For experiments involving constant illumination and 12 hour light dark cycle, incubators were maintained at 25°C with a white light source (light intensity: 8000 lux) for the required time period. For constant dark rearing, flies within the vials were kept in tightly closed dark boxes and maintained in the constant temperature laboratory incubator. Gal4-UAS system was used for the selective activation of the transgene spatially and temporally for targeted gene expression.

### Sequence and Phylogeny Analysis

The protein sequences for sequence alignment were obtained from NCBI: dEsyt-PA (NP_733010.1), hEsyt1- isoform 1 (NP_001171725.1), hEsyt2- isoform 1 (NP_001354702.1), hEsyt3- isoform a (NP_001309760.1), yeast tcb1 (NP_014729.1), yeast tcb2 (NP_014312.1), yeast tcb3 (NP_013639.1). Multiple sequence alignment of dEsyt protein sequence with the human Esyts and yeast tricalbins were obtained using MUSCLE (MUltiple Sequence Comparison by Log-Expectation) (Edgar, 2004). The phylogenetic tree reflecting the evolution of these proteins were obtained using neighbour joining method MEGA6 (Molecular Evolutionary Genetics Analysis Version 6.0) (Tamura et al., 2013).

### Optical Neutralization

The flies reared under experimental conditions were immobilized by cooling on ice, carefully decapitated and fixed on the microscope slide using a drop of colourless nail varnish. The refractive index of the cornea was neutralized using a drop of immersion oil, viewed and imaged under the 40X oil immersion objective of Olympus BX43 microscope. The digital image acquisition and documentation was done by using CellSens software.

### Scoring retinal degeneration

To obtain a quantitative index of degeneration, a total of 50 ommatidia from 5 different flies of each genotype were assessed for each time point. The single, central UV sensitive photoreceptor that didn’t show any light dependent retinal degeneration was used as the reference and the rest of the photoreceptors were scored. A score of 1 was given to each rhabdomere that appeared to be wild type. Thus the control photoreceptors will have a score of 7 and the mutants undergoing degeneration will have a score from 1 to 7. The results were analysed and the graph was plotted using GraphPad Prism software.

### Electroretinogram

Anesthetized flies were introduced into truncated 200 μl disposable pipette tips such that the head protruded from the small opening. The fly’s head was then immobilized using colourless nail varnish. Two glass microelectrodes (640786, Harvard Apparatus, Massachusetts, USA) filled with 0.8% NaCl solution were used for recordings such that the voltage changes were recorded by placing the experimental electrode on the surface of the eye and the reference electrode on the thorax. The flies were dark adapted for 5 minutes prior to ERG recordings followed by 2 sec flash of green light stimulus, with 10 stimuli (flashes) per recording and 15 seconds of recovery time between two subsequent flashes. Green light stimulus was emitted using an LED light source to within 5 mm of fly’s eye through a fibre optic guide. Voltage changes were recorded using pCLAMP 10.7 and amplified using DAM50 amplifier (SYS-DAM50, WPI, Florida, USA). Data analysis was done using Clampfit 10.7 (Molecular Devices, California, USA). Graphs were plotted using GraphPad Prism software.

### Western blot

Age matched flies were decapitated and the heads were homogenized in 2X laemmli sample buffer followed by boiling at 95°C for 5 minutes. For rhodopsin blot, the protein samples were processed differently. The fly heads were snap frozen in liquid nitrogen and stored at −80°C for a period of two days. These head samples were homogenised in 2X laemmli sample buffer followed by boiling at 37°C for 30 minutes. Protein extracts from fly heads were separated using SDS-PAGE and transferred onto nitrocellulose filter membrane [Hybond-C Extra; (GE Healthcare, Buckinghamshire, UK)] using wet transfer apparatus (BioRad, California, USA). The membrane was blocked using 5% Blotto (sc-2325, Santa Cruz Biotechnology, Texas, USA) in Phosphate Buffer Saline (PBS) with 0.1% Tween 20 (Sigma Aldrich) (PBST) for 2 hours at room temperature (RT). Primary antibody incubation was done overnight at 4⁰C using appropriate antibody dilutions: anti-RDGB (lab generated), 1:1000; anti-dEsyt (kind gift from Dr. Dion Dickman’s lab), 1:2000; anti-rhodopsin (4C5-DHSB, lowa, USA), 1:250; anti-Gq, 1:1000; anti-TRP (lab generated), 1:4000; anti-norpA,1:1000 and anti-α-tubulin-E7c (DHSB, lowa, USA), 1:4000. Following this, the membrane was washed in PBST and incubated with 1:10000 dilutions of appropriate secondary antibody (Jackson ImmunoResearch Laboratories, Pennsylvania, USA) coupled to horseradish peroidase at RT for 2 hours. Three PBST washes were given and the blots were developed with ECL (GE Healthcare) and imaged using LAS 4000 instrument (GE Healthcare).

### Fluorescent pseudopupil

To monitor changes in PIP_2_ levels at the microvillar PM in live flies, the PIP_2_ biosensor PH-PLCδ coupled to GFP driven by the transient receptor (*trp*) promoter of flies was used (Yadav et al., 2015). The flies were made insentient and were immobilised using the same protocol as in ERG recordings. The pseudopupil formed from the summed fluorescence of approximately 20-40 adjacent ommatidia were focused and imaged using the 10X objective of Olympus IX71 microscope. The program created using the software Micromanager captured time lapse images of the pseudopupil by collecting fluorescence emitted from the eye when GFP was stimulated by a 90 ms flash of blue light. Prior to the recordings, the flies were dark adapted (resting conditions) for 6 minutes during which the probe binds to PIP_2_ and localizes to the microvillar PM. Following a flash of blue light, the PLCβ activity triggers the hydrolysis of PIP_2_ and thereby the probe is displaced from the PM and this leads to the loss of the pseudopupil. The central UV sensitive photoreceptor is unresponsive to blue light and thereby retains the probe at the apical PM. Subsequent red light illuminations following the blue light stimulus after each time lapse image acquisition hastens the retrieval of the probe back to the PM. Mean fluorescence intensity indicating the basal PIP_2_ pools and the PIP_2_ recovery kinetics were calculated using ImageJ from NIH (Bethesda, MD, USA). Quantification of DPP fluorescence intensity was done by measuring the intensity values per area of the pseudopupil.

Similarly, flies expressing the P4M-GFP probe (PI4P biosensor) were subjected to the same protocol for monitoring PI4P levels. P4M-GFP expression was driven by an eye specific promoter GMR using Gal4-UAS system.

### RNA extraction and qPCR

RNA isolation was done from *Drosophila* heads using TRIzol reagent (Invitrogen) followed by treatment with amplification grade DNAse I (Invitrogen). cDNA synthesis was done using the Superscript II RNAse H Reverse Transcriptase (Invitrogen) and random hexamers (Applied biosystems). dEsyt primers were designed at the exon-exon junction of the dEsyt genomic region satisfying the parameters recommended for q-PCR primer designing and the run was performed in the Applied Biosystem 7500 Fast Real Time PCR instrument using Ribosomal Protein 49 (RP49) primers as the control primers flanking the house keeping gene region. Triplicates of each sample were measured to ensure the consistency of the data. The primers used for q-PCR include:

RP49 forward: CGGATCGATATGCTAAGCTGT
RP49 reverse: GCGCTTGTTCGATCCGTA
dEsyt forward: GTTGTGGATAGTTGGCTCACCTT
dEsyt reverse: GCCTGGCTGAATCGATGAATAC
U6.1 forward: GATCCTGTGGCGGCTAC
U6.1 reverse: GAGGTGGAGTCTGGAAAGC

### Immunohistochemistry

Retinae were dissected in PBS and fixed with 4% paraformaldehyde in PBS with 1 mg/ml saponin at room temperature for 30 min. Post fixation, the samples were washed in PBS with 0.3% Triton X-100 (0.3% PBTX) followed by incubation with blocking solution (5% Fetal Bovine solution in PBTX) for 2 hours at RT. The samples were then incubated overnight with the respective antibody [anti-RDGB (lab generated); 1:300, dEsyt (kind gift from Dr. Dion Dickman’s lab); 1:200, mCherry (ThermoFisher Scientific-PA5-34974); 1:300] at 4°C. Post this, samples were washed thrice with 0.3% PBTX and incubated with appropriate secondary antibody [Alexa Fluor 488 anti-rabbit (A11034), anti-rat (A11006) and Alexa Fluor 633 anti-rabbit (A21070), Molecular Probes; 1:300 dilution] for 4 hours at RT. Along with the secondary antibody incubation, Alexa Fluor 568–Phalloidin (Invitrogen, A12380; 1:300) was used to mark F-actin. Samples were washed thrice with 0.3% PBTX, followed by one final wash in PBS and were mounted with 70% glycerol in 1X PBS. The whole-mounted preparations were imaged under 60X 1.4 NA objective, in Olympus FV 3000 microscope.

### Electron Microscopy

The fly heads of mentioned genotypes were cut and immersed in 2% osmium tetroxide, kept at 4°C for 1 hour followed by incubation at 40°C for 4 days. Specimens were washed with distilled water, stained *enbloc* with uranyl acetate (0.5% in distilled water) for 3 hours. After washing with distilled water, specimens were subjected to dehydration step and embedded in epon. Ultrathin sections of 60 nm were cut and grids were subjected to post staining with 2% uranyl acetate (in 70% ethanol) and Reynold’s lead solution. Sections were imaged at 120 KV on a Tecnai G2 Spirit Bio-TWIN (FEI) electron microscope.

### Scoring MCS density

For scoring MCS density/photoreceptor cell, a total of 30 cells were taken to conduct analysis for R1-R6 and 9 cells for R7. Using freehand line tool of image J, length of MCS (μm) /the total length of the base of the rhabdomere (μm) was calculated. Fractions of MCS coverage were multiplied with 100 to show the percentage. Groups were compared using the GraphPad Prism 8 software.

### Statistical analysis

Unpaired two tailed t test or ANOVA, followed by Tukey’s multiple comparison test, were carried out where applicable.

## Supplementary figure legends

**Supplementary figure 1: Reduced mass and body size in *dEsyt*^*KO*^ adults**

**(A)** Representative image of adult flies showing the reduced body size in *dEsyt* knockout mutants.

**(B)** Quantification of the reduced mass of *dEsyt*^*KO*^ v Wild type. n=10 flies.

**(C)** Representative image showing the rescue of body size by the transgenic expression of dEsyt∷mCherry in *dEsyt*^*KO*^ homozygotes.

**(D)** Quantification of the rescue in body mass of flies shown in image C. n=10 flies

Scatter plots with mean ± SD are shown. Statistical tests: (B) Student’s unpaired t test. (D) one-way ANOVA with post hoc Tukey’s multiple pairwise comparison. ***p value <0.001; ****p value <0.0001.

**Supplementary Figure 2: Loss of *dEsyt* delays the PI4P recovery kinetics in *rdgB*^*9*^.**

**(A, C)** Quantification of the mean fluorescence intensity of the deep pseudopupil formed by the PI4P probe P4M-GFP in one day old flies of the indicated genotypes (n=10).

**(B, D)** Graph showing the recovery kinetics of the fluorescent pseudopupil with time, X-axis represent the genotypes, Y-axis represent intensity expressed as percentage intensity of each pseudopupil acquired at different time points normalized to the intensity of the first image acquired (n=10).

(E) Representative deep pseudopupil images from one day old flies expressing P4M-GFP probe acquired at specified time points. Genotypes as indicated (n=10).

Scatter plots and XY plots with mean ± SD are shown. Statistical tests: (A and C) Student’s unpaired t test. (B and D) Two-Way ANOVA Grouped analysis with Bonferroni post-tests to compare replicate means. ns- Not significant; **p value < 0.01; ***p value < 0.001; ****p value <0.0001.

**Supplementary Figure 3: Loss of *dEsyt* does not affect the levels of key phototransduction proteins in *rdgB^9^***

Western blot of head extracts made from flies of the indicated genotypes. Tubulin was used as the loading control for the experiment. The blots were probed for the major phototransduction proteins **(A)** Rhodopsin **(B)** Gq **(C)** PLC **(D)** TRP.

**Supplementary Figure 4: ERG amplitude remains unaffected in *dEsyt* knocked out *PI4KIIIα* mutants.**

**(A)** Average ERG response of 0-1 day dark reared flies of the indicated genotype to a 2s flash of green light; n=10.

**(B)** Representative traces of ERG showing the duration of light pulse, X-axis indicates time in msec and Y-axis indicates the average ERG amplitude in mV.

Scatter plots with mean ± SD are shown. Statistical tests: (A) one-way ANOVA with post hoc Tukey’s multiple pairwise comparison. ns- Not significant.

## References

Balakrishnan, S.S., Basu, U., Shinde, D., Thakur, R., Jaiswal, M., and Raghu, P. (2018). Regulation of PI4P levels by PI4KIIIα during G-protein-coupled PLC signaling in *Drosophila* photoreceptors. J. Cell Sci. 131, jcs217257.

Balla, T., Kim, Y.J., Alvarez-Prats, A., and Pemberton, J. (2019). Lipid Dynamics at Contact Sites Between the Endoplasmic Reticulum and Other Organelles. Annu. Rev. Cell Dev. Biol. 35, 85–109.

Bian, X., Saheki, Y., and Camilli, P. De (2018). ER-plasma membrane tethering with lipid transport. EMBO J 37, 219–234.

Carrasco, S., and Meyer, T. (2014). NIH Public Access. 973–1000.

Chang, C.-L., Hsieh, T.-S., Yang, T.T., Rothberg, K.G., Azizoglu, D.B., Volk, E., Liao, J.-C., and Liou, J. (2013). Feedback regulation of receptor-induced Ca2+ signaling mediated by E-Syt1 and Nir2 at endoplasmic reticulum-plasma membrane junctions. Cell Rep. 5, 813–825.

Cockcroft, S., and Raghu, P. (2018). Phospholipid transport protein function at organelle contact sites. Curr. Opin. Cell Biol. 53, 52–60.

Cockcroft, S., Garner, K., Yadav, S., Gomez-Espinoza, E., and Raghu, P. (2016). RdgBα reciprocally transfers PA and PI at ER-PM contact sites to maintain PI(4,5)P2 homoeostasis during phospholipase C signalling in Drosophila photoreceptors. Biochem. Soc. Trans. 44, 286–292.

Edgar, R.C. (2004). MUSCLE : multiple sequence alignment with high accuracy and high throughput. Nucleic Acids Res. 32, 1792–1797.

Gatta, A.T., and Levine, T.P. (2017). Piecing Together the Patchwork of Contact Sites. Trends Cell Biol. 27, 214–229.

Giordano, F., Saheki, Y., Idevall-Hagren, O., Colombo, S.F., Pirruccello, M., Milosevic, I., Gracheva, E.O., Bagriantsev, S.N., Borgese, N., and De Camilli, P. (2013). PI(4,5)P(2)-dependent and Ca(2+)-regulated ER-PM interactions mediated by the extended synaptotagmins. Cell 153, 1494–1509.

Hayashi, M., Raimondi, A., Toole, E.O., Paradise, S., Collesi, C., Cremona, O., Ferguson, S.M., and Camilli, P. De (2008). Cell- and stimulus-dependent heterogeneity of synaptic vesicle endocytic recycling mechanisms revealed by studies of dynamin 1-null neurons. Proc Natl Acad Sci USA 105, 1–6.

Hepler, P.K., Palevitz, B.A., Lancelle, S.A., Mccauley, M.M., and Lichtscheidl, I. (1984). Cortical endoplasmic reticulum in plants. J Cell Sci. 96, 355–373.

Herdman, C., Tremblay, M.G., Mishra, P.K., and Moss, T. (2014). Loss of Extended Synaptotagmins ESyt2 and ESyt3 does not affect mouse development or viability, but in vitro cell migration and survival under stress are affected. Cell Cycle 13, 2616–2625.

Idevall-hagren, O., Lü, A., Xie, B., and Camilli, P. De (2015). Triggered Ca 2 + influx is required for extended membrane tethering. EMBO J 34, 2291–2305.

Kikuma, K., Li, X., Kim, D., Sutter, D., and Dickman, D.K. (2017). Extended Synaptotagmin Localizes to Presynaptic ER and Promotes Neurotransmission and Synaptic Growth in Drosophila. Genetics 207, 993–1006.

Liu, W., Xie, Y., Ma, J., Luo, X., Nie, P., Zuo, Z., Lahrmann, U., Zhao, Q., Zheng, Y., Zhao, Y., et al. (2015). IBS: an illustrator for the presentation and visualization of biological sequences. Bioinformatics 31, 3359–3361.

Loewen, C.J.R., Young, B.P., Tavassoli, S., and Levine, T.P. (2007). Inheritance of cortical ER in yeast is required for normal septin organization. J Cell Biol 179, 467–483.

Manford, A.G., Stefan, C.J., Yuan, H.L., MacGurn, J.A., and Emr, S.D. (2012). ER-to-Plasma Membrane Tethering Proteins Regulate Cell Signaling and ER Morphology. Dev. Cell 23, 1129–1140.

Omnus, D.J., Manford, A.G., Bader, J.M., Emr, S.D., and Stefan, C.J. (2016). Phosphoinositide kinase signaling controls ER-PM cross-talk. Mol. Biol. Cell 27, 1170–1180.

Palade, G.E., and Porter, K.R. (1957). Studies on the endoplasmic reticulum: Its form and distribution in striated muscle cells. J Biophys.Biochem.Cytol 3, 269–300.

Pearn, M.T., Randall, L.L., Shortridge, R.D., Burg, M.G., and Pak, W.L. (1996). Molecular, biochemical, and electrophysiological characterization of Drosophila norpA mutants. J Biol Chem 271, 4937–4945.

Pichler, H., Gaigg, B., Hrastnik, C., Achleitner, G., Kohlwein, S.D., and Daum, È. (2001). A subfraction of the yeast endoplasmic reticulum associates with the plasma membrane and has a high capacity to synthesize lipids. Eur J Biochem 268, 2351–2361.

Raghu, P., Yadav, S., and Mallampati, N.B.N. (2012). Lipid signaling in Drosophila photoreceptors. Biochim. Biophys. Acta 1821, 1154–1165.

Saheki, Y., and De Camilli, P. (2017). Endoplasmic Reticulum–Plasma Membrane Contact Sites. Annu. Rev. Biochem. 86, 659–684.

Saheki, Y., Bian, X., Schauder, C.M., Sawaki, Y., Surma, M.A., Klose, C., Pincet, F., Reinisch, K.M., and De Camilli, P. (2016). Control of plasma membrane lipid homeostasis by the extended synaptotagmins. Nat. Cell Biol. 18, 504–515.

Schauder, C.M., Wu, X., Saheki, Y., Narayanaswamy, P., Torta, F., Wenk, M.R., De Camilli, P., and Reinisch, K.M. (2014). Structure of a lipid-bound extended synaptotagmin indicates a role in lipid transfer. Nature 510, 552–555.

Sclip, A., Bacaj, T., Giam, L.R., and Südhof, T.C. (2016). Extended Synaptotagmin (ESyt) Triple Knock- Out Mice Are Viable and Fertile without Obvious Endoplasmic Reticulum Dysfunction. PLoS One 11, 1–17.

Stefan, C.J., Manford, A.G., Baird, D., Yamada-Hanff, J., Mao, Y., and Emr, S.D. (2011). Osh proteins regulate phosphoinositide metabolism at ER-plasma membrane contact sites. Cell 144, 389–401.

Suzuki, E., and Hirosawa, K. (1994). Immunolocalization of a Drosophila Phosphatidylinositol Transfer Protein (rdgB) in Normal and rdgA Mutant Photoreceptor Cells with Special Reference to the Subrhabdomeric Cisternae. J. Electron. Microsco. 189, 183–189.

Tamura, K., Stecher, G., Peterson, D., Filipski, A., and Kumar, S. (2013). MEGA6: Molecular Evolutionary Genetics Analysis version 6.0. Mol. Biol. Evol. 30, 2725–2729.

Tremblay, M.G., and Moss, T. (2016). Loss of all 3 Extended Synaptotagmins does not affect normal mouse development, viability or fertility. Cell Cycle 15, 2360–2366.

Vihtelic, T.S., Goebl, M., Milligan, S., O’Tousa, J.E., and Hyde, D.R. (1993). Localization of Drosophila retinal degeneration B, a membrane-associated phosphatidylinositol transfer protein. J. Cell Biol. 122, 1013–1022.

Yadav, S., Garner, K., Georgiev, P., Li, M., Gomez-Espinosa, E., Panda, A., Mathre, S., Okkenhaug, H., Cockcroft, S., and Raghu, P. (2015). RDGBα, a PtdIns-PtdOH transfer protein, regulates G-proteincoupled PtdIns(4,5)P2 signalling during Drosophila phototransduction. J. Cell Sci. 128, 3330–3344.

Yadav, S., Cockcroft, S., and Raghu, P. (2016). The Drosophila photoreceptor as a model system for studying signalling at membrane contact sites. Biochem. Soc. Trans. 44, 447–451.

Yadav, S., Thakur, R., Georgiev, P., Deivasigamani, S., K, H., Ratnaparkhi, G., and Raghu, P. (2018). RDGBα localization and function at a membrane contact site is regulated by FFAT/VAP interactions. J. Cell Sci. 131, doi:10.1242/jcs.207985.

Yu, H., Liu, Y., Gulbranson, D.R., Paine, A., Rathore, S.S., and Shen, J. (2016). Extended synaptotagmins are Ca ^2+^ -dependent lipid transfer proteins at membrane contact sites. Proc. Natl. Acad. Sci. 113, 4362–4367.

