## Supplementary material for "Extended synaptotagmin regulates plasma membrane-endoplasmic reticulum contact site structure and lipid transfer function *in vivo*": Esyt_Sup fig

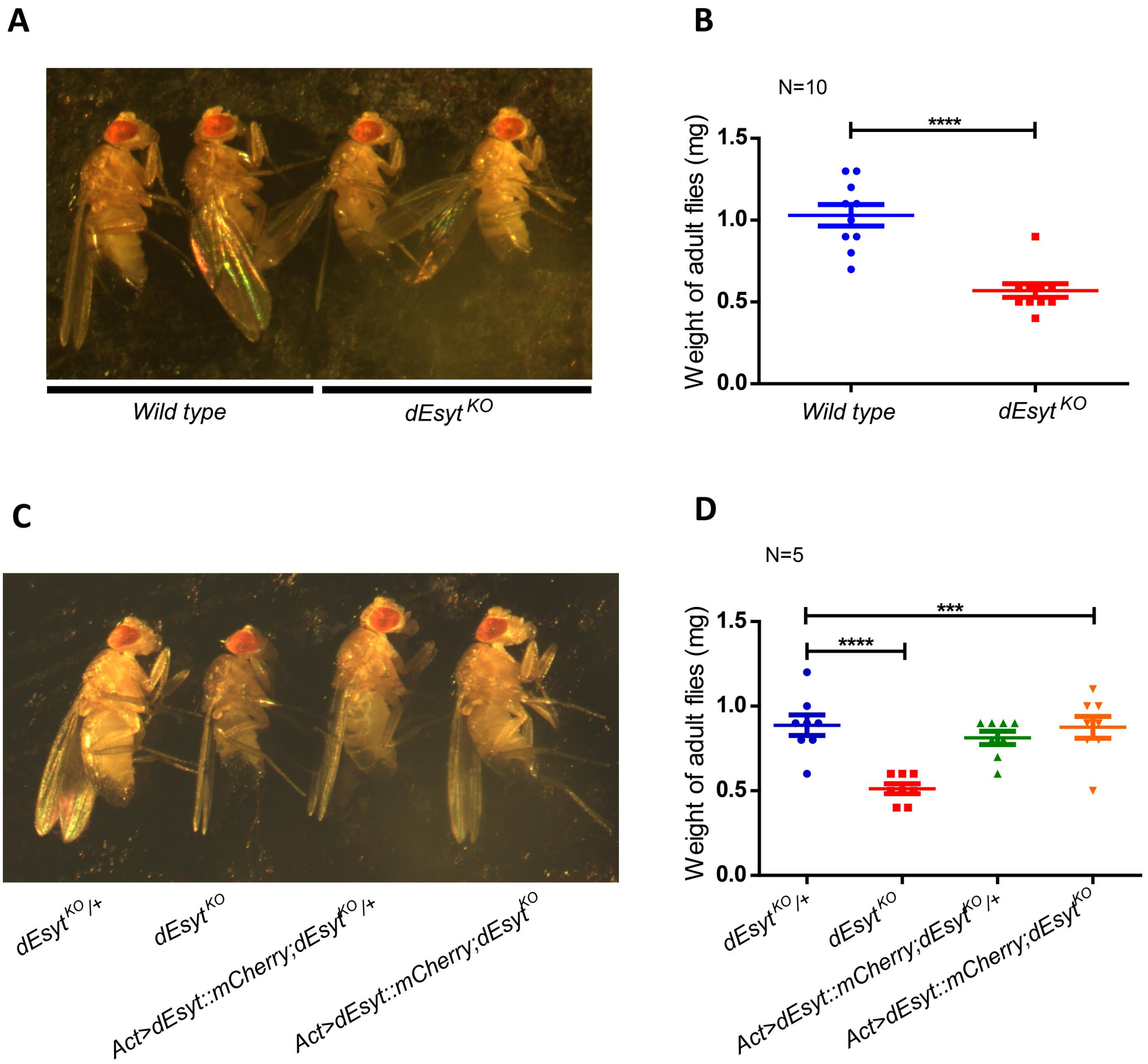


**Nath et. al, 2019 Supplementary Figure 1**

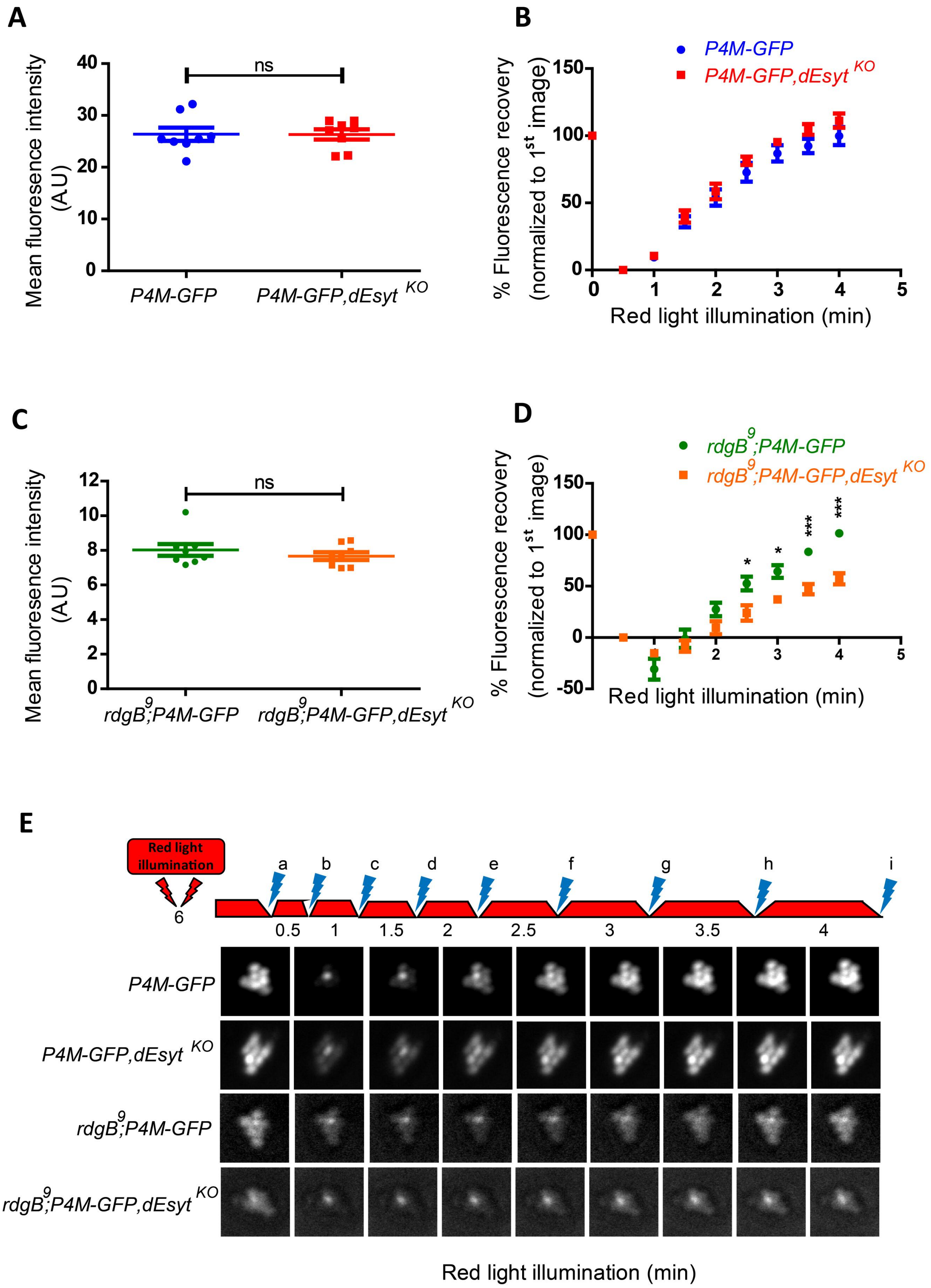


**Nath et. al, 2019 Supplementary Figure 2**

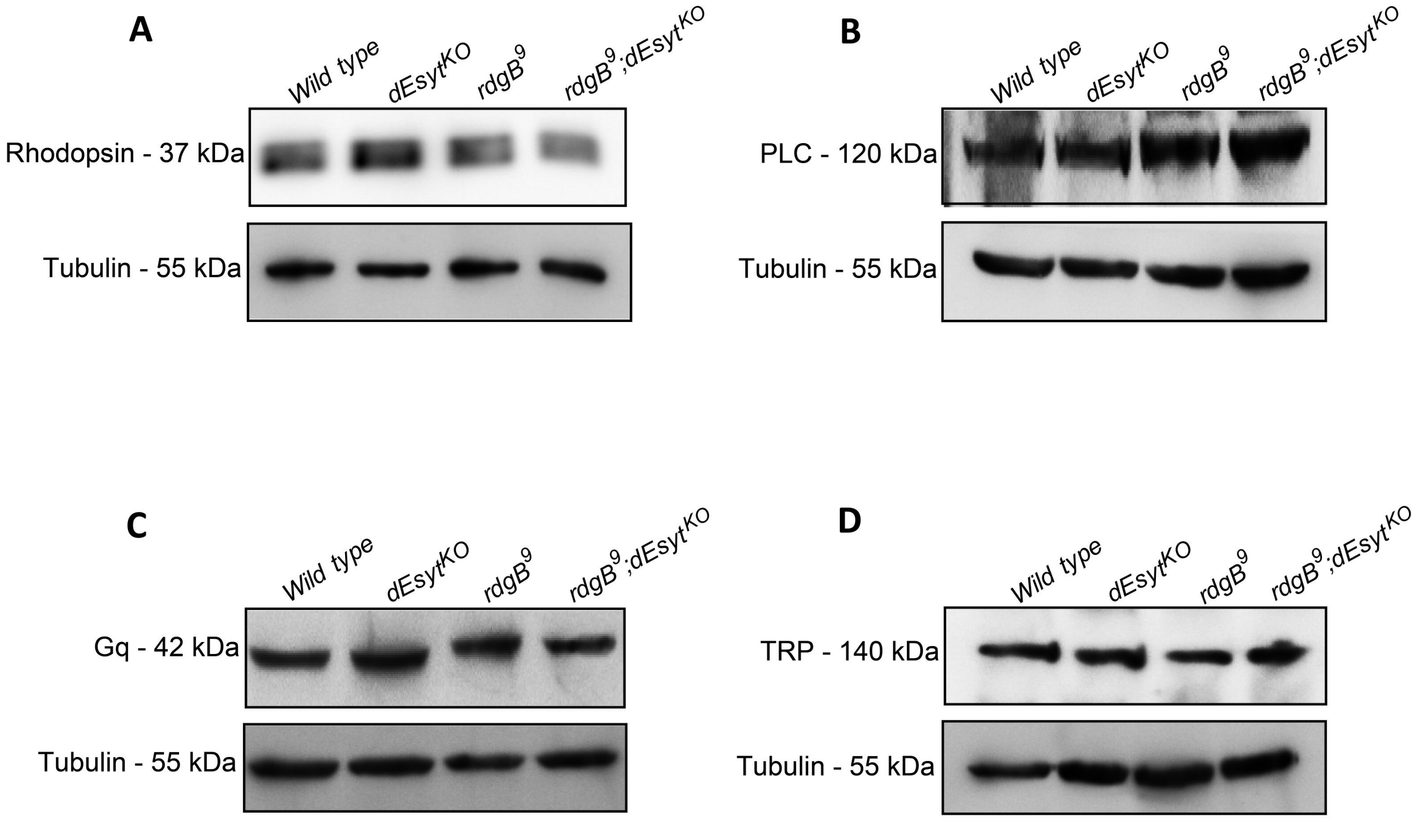


**Nath et. al, 2019 Supplementary Figure 3**

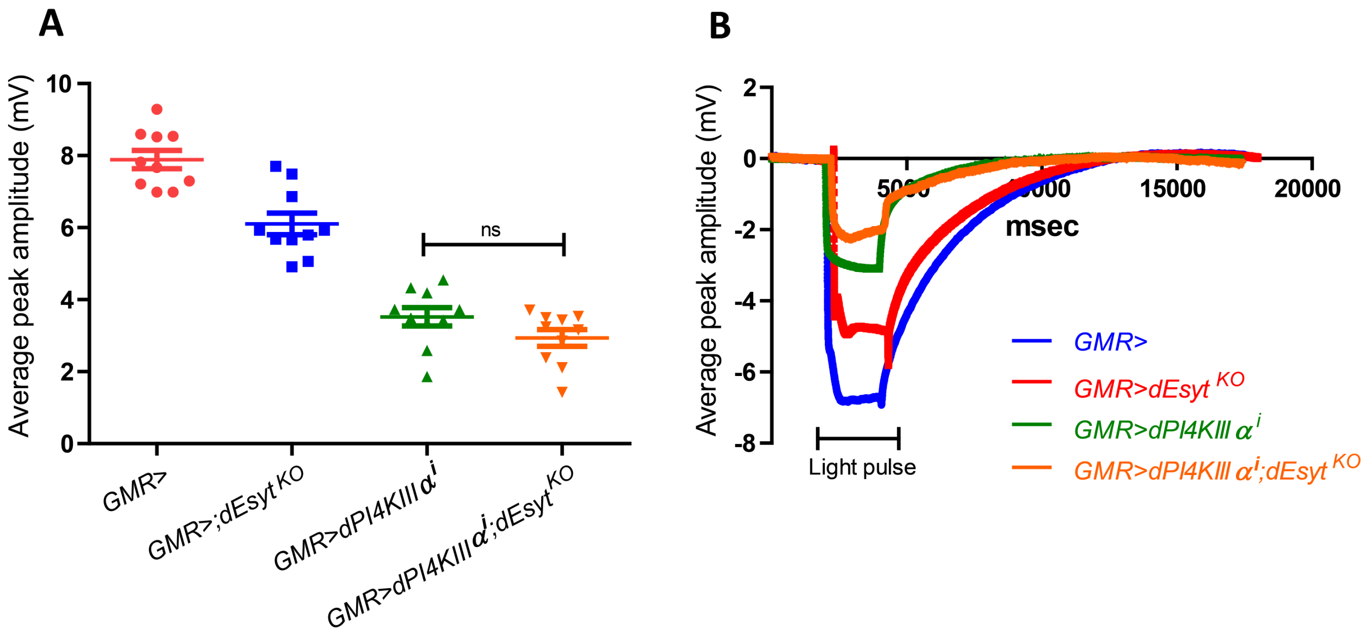


**Nath et. al, 2019 Supplementary Figure 4**
